# A model for accurate quantification of CRISPR effects in pooled FACS screens

**DOI:** 10.1101/2024.06.17.599448

**Authors:** Harold Pimentel, Jacob W. Freimer, Maya M. Arce, Christian M. Garrido, Alexander Marson, Jonathan K. Pritchard

## Abstract

CRISPR screens are powerful tools to identify key genes that underlie biological processes. One important type of screen uses fluorescence activated cell sorting (FACS) to sort perturbed cells into bins based on the expression level of marker genes, followed by guide RNA (gRNA) sequencing. Analysis of these data presents several statistical challenges due to multiple factors including the discrete nature of the bins and typically small numbers of replicate experiments. To address these challenges, we developed a robust and powerful Bayesian random effects model and software package called Waterbear. Furthermore, we used Waterbear to explore how various experimental design parameters affect statistical power to establish principled guidelines for future screens. Finally, we experimentally validated our experimental design model findings that, when using Waterbear for analysis, high power is maintained even at low cell coverage and a high multiplicity of infection. We anticipate that Waterbear will be of broad utility for analyzing FACS-based CRISPR screens.

## Introduction

Genetic screening is a powerful technique to identify the genes that underlie a phenotype or that are involved in a particular biological process. The ability of CRISPR/Cas9 to induce genetic perturbations efficiently has facilitated large-scale screens in many mammalian cell types ^1,2^. CRISPR screens can be paired with FACS to map the genetic wiring underlying complex phenotypes by identifying key upstream regulators of specific, relevant target genes ^3–8^. These screens use fluorescent reporters or fluorescent antibodies to directly measure the expression level of a protein of interest or a protein that is a surrogate marker of a biological process (such as a phosphorylated protein at the end of a signaling cascade). For simplicity, we will refer to any target measured by FACS – either an endogenous protein or a reporter protein – as a marker throughout the rest of the text. After pooled CRISPR perturbations, FACS is used to sort cells into different bins based on the fluorescence intensity of the marker. By sequencing the relative abundance of gRNAs in each bin it is possible to associate genetic perturbations with their effect on the levels of the marker.

There are a number of experimental and computational challenges when performing CRISPR FACS screens. These screens must balance a desire to perturb many genes, with high cell coverage for each perturbation, against costs and experimental demands. Furthermore, there is increasing interest in performing such screens in primary cells or in *in vivo* models which are more relevant for disease, but for which the number of cells that can be used is often limited ^9–11^. CRISPR screens are also usually only performed with two or three replicates. Limiting numbers of cells and replicates reduce the number of times each gRNA is measured, which increases noise and uncertainty. This noise compounds with other sources of variability between replicates and donors, between different gRNAs targeting the same gene, and imprecise FACS gates.

Finally, FACS screens are further complicated by the fact that they involve a pool of perturbed cells so the effect of each gRNA on the marker cannot be measured directly, but must instead be inferred by the relative abundance of gRNAs in different FACS bins. These challenges necessitate the development of new analysis methods designed specifically to analyze CRISPR FACS screens. Furthermore, there are often not principled guidelines on how different parameters affect the statistical power of these screens to inform experimental design.

We developed a computational framework, Waterbear, that (1) performs robust inference of CRISPR FACS screens and (2) informs optimal experimental design by iterating over thousands of plausible experimental configurations through simulation. Given parameters learned from real data, the model can generate realistic simulations of experiments at the single-cell level using a generative view of the model. The generative view of the model enables stepping through each parameter of the model to generate data that is consistent with parameters learned from real data while still introducing randomness at every stage of the model consistent with biological and experimental variability. Once a simulation is done, the statistical power of that experimental configuration can be estimated using our gene-level inference model which is a simplified version of the cell-level model which more closely mimics how the data is observed in practice. The inference model aggregates the cell-level information into a count observation and models the gRNA count distribution across discrete bins as is observed in actual screens. Waterbear is robust in that it can infer bin sizes, model the latent effects of the gRNAs on the marker distribution, and share information across guides, genes, and replicates to assess uncertainty. Further, this model is also used to analyze real data where the cell-level information is not available.

Waterbear is designed to use all available information to make informed decisions about whether each perturbed gene affects the marker distribution by modeling several relationships that are inherent to such screens. This model enables inferring thousands of parameters by shrinking the results towards a shared prior across relevant dimensions. For example, on average, gRNAs targeting the same gene should behave similarly, so in the model, gRNAs targeting the same gene share a “parent” distribution, while allowing each guide to have a unique effect size. While some gRNAs will produce off-target effects, Waterbear’s design does not ignore them, but rather downweights the evidence of the gene-level effect size if the off-target gRNA is inconsistent with other guides. Similarly, negative controls are used to infer experiment-level parameters such as the null marker distribution, variance between replicates, and variance between gRNAs. Additionally, Waterbear uses a sparse prior for *gene-level* effects since out of thousands of gRNAs, only a modest fraction of gRNAs will have a true measurable effect on the marker distribution.

While other tools have been used widely for FACS-based screen analysis, the current tools all have limitations for this purpose. MAGeCK was originally designed to analyze cell abundance screens and is one of the most commonly used CRISPR screen analysis tools ^12^. However, MAGeCK only supports comparisons between two populations, and therefore it cannot take advantage of the additional information collected with more than two FACS bins, preventing it from modeling the underlying marker distribution. In contrast, MAUDE was developed specifically for the analysis of FACS screens ^13^. However, MAUDE does not explicitly handle replicates, requires a separate input population, and requires precise bin sizes to be manually specified. Finally, RELICS shares technical similarities with Waterbear as it is also a Bayesian hierarchical model. However, RELICS is designed for CRISPR tiling screens that perturb non-coding sequences where one would expect spatial correlations between guides, and thus the software is designed around the concept of finding “functional sites’’, rather than finding gene-gene relationships as we describe here. Waterbear was designed to overcome these limitations.

In addition to being able to analyze data, an expanded cell-level view of the Waterbear model can be used to simulate experiments to explore a wide range of experimental parameters and their implication on the design of these screens. We used Waterbear to explore how FACS bin size, gRNA coverage (the number of times a gRNA is measured), and lentiviral gRNA library multiplicity of infection ^14^ (MOI) affect the power of FACS-based screens and show that it is possible to reduce the number of cells required while still maintaining high sensitivity. Unlike existing simulation methods ^15^, we simulate each cell individually, enabling us to change cell-level parameters such as the MOI. We validate our simulations and inference procedure by repeating previous screens ^3^ at a higher MOI and with lower coverage of the gRNAs. Our results provide a roadmap to reduce the number of cells required for FACS screens and we introduce a powerful new analysis tool to analyze such screens. These advances open the door for future screens addressing novel biological questions in rare, primary cell types and *in vivo* models.

Waterbear is available as free and open-source software at https://github.com/pimentel/waterbear.

## Results

### Overview of CRISPR FACS screens and tunable parameters

To understand the genetic regulation of a marker of interest, the most straightforward approach would be to individually perturb the expression of candidate regulatory genes and then measure the expression of the marker. This approach is challenging to scale and instead perturbations are often performed in a pool of cells with each cell containing a different perturbation ^16^. When the cells are pooled, the distribution of the marker reflects a mixture of many different perturbations and the effect of each individual perturbation cannot be directly observed (Figure 1A). However, using FACS, the perturbed cells can be sorted into different bins based on the expression of the marker ^16^. By sequencing the gRNAs in the sorted cells and identifying which gRNAs are differentially enriched between the sorted populations, one can identify regulators of the marker (Figure 1A).

**Figure 1:**
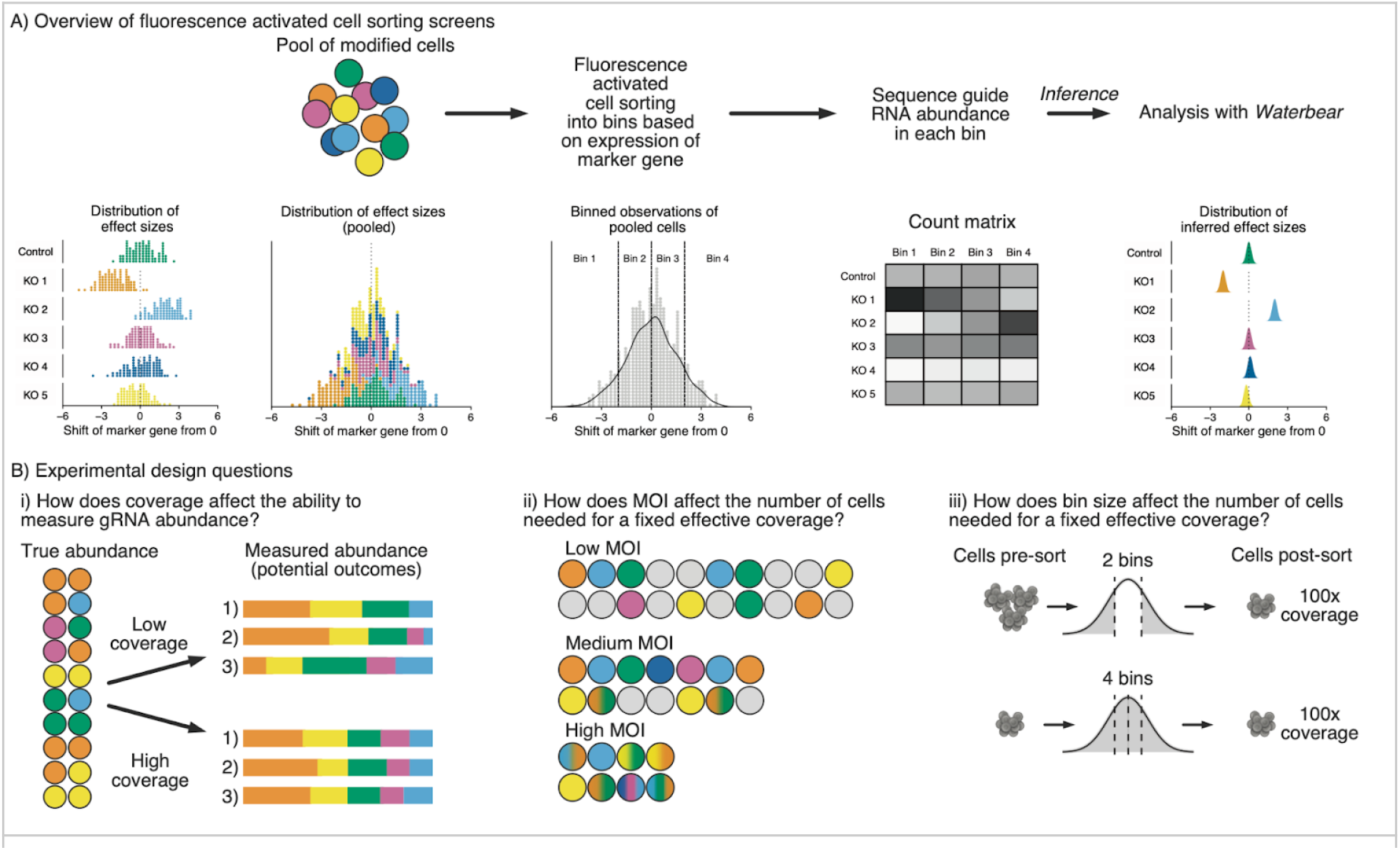
Overview of CRISPR FACS screens and tunable parameters. A) Schematic of pooled CRISPR FACS screens. Perturbations that differentially affect the expression of a target protein of interest are mixed together in a pool of modified cells. To infer the effect of each gRNA, cells are sorted into bins based on expression of a target protein using FACS; gRNA abundance is compared across bins through sequencing. B) Experimental design considerations focused on reducing cell requirements, including the effect of changing i. gRNA coverage, ii. MOI, and iii. FACS bin configuration.

The field lacks principled guidelines for how experimental design choices affect the statistical power to detect differentially enriched gRNAs. Coverage, MOI, and FACS bin size are highly interrelated experimental parameters that can be altered to balance the number of cells needed for an experiment and the accuracy of gRNA abundance measurements (Figure 1B). For instance, at high MOI there will be multiple gRNAs per cell, which increases the effective coverage for a fixed number of cells while also increasing the variance of the observations. Here, we provide a framework to explore how these experimental parameters affect false discoveries and statistical power and provide principled guidelines for future screens.

### A statistical model for CRISPR FACS screen data

We present two possible generative views of the data; one with cell-level information (the “cell-level” view) and one with aggregate information as is observed in FACS screen data (the “gene-level” view). The cell-level view offers details on all guides within a single cell (as in in single-cell sequencing), while the gene-level view provides only the total occurrences of a guide-bin combination, the default in FACS-based screens. For simplicity, we describe the gene-level model in detail below, but the more general cell-level model is discussed in detail in Supplementary Section 1. Importantly, we perform simulated screens using the cell-level model while iterating over the parameter space, but perform inference using the gene-level model to match the data available from a typical FACS screen.

In the Waterbear model (Figure 2A), there are two classes of genes, those that have no effect on the marker (denoted by *ψ_i_* = 0), and those that have an effect on the marker (*ψ_i_* = 1). By classifying genes into two discrete groups, our model defines different gRNA-marker behavior based on the inferred class. The central goal of Waterbear is to report, for each gene, the posterior probability that *ψ_i_* = 1, as well as estimated effect sizes.

**Figure 2:**
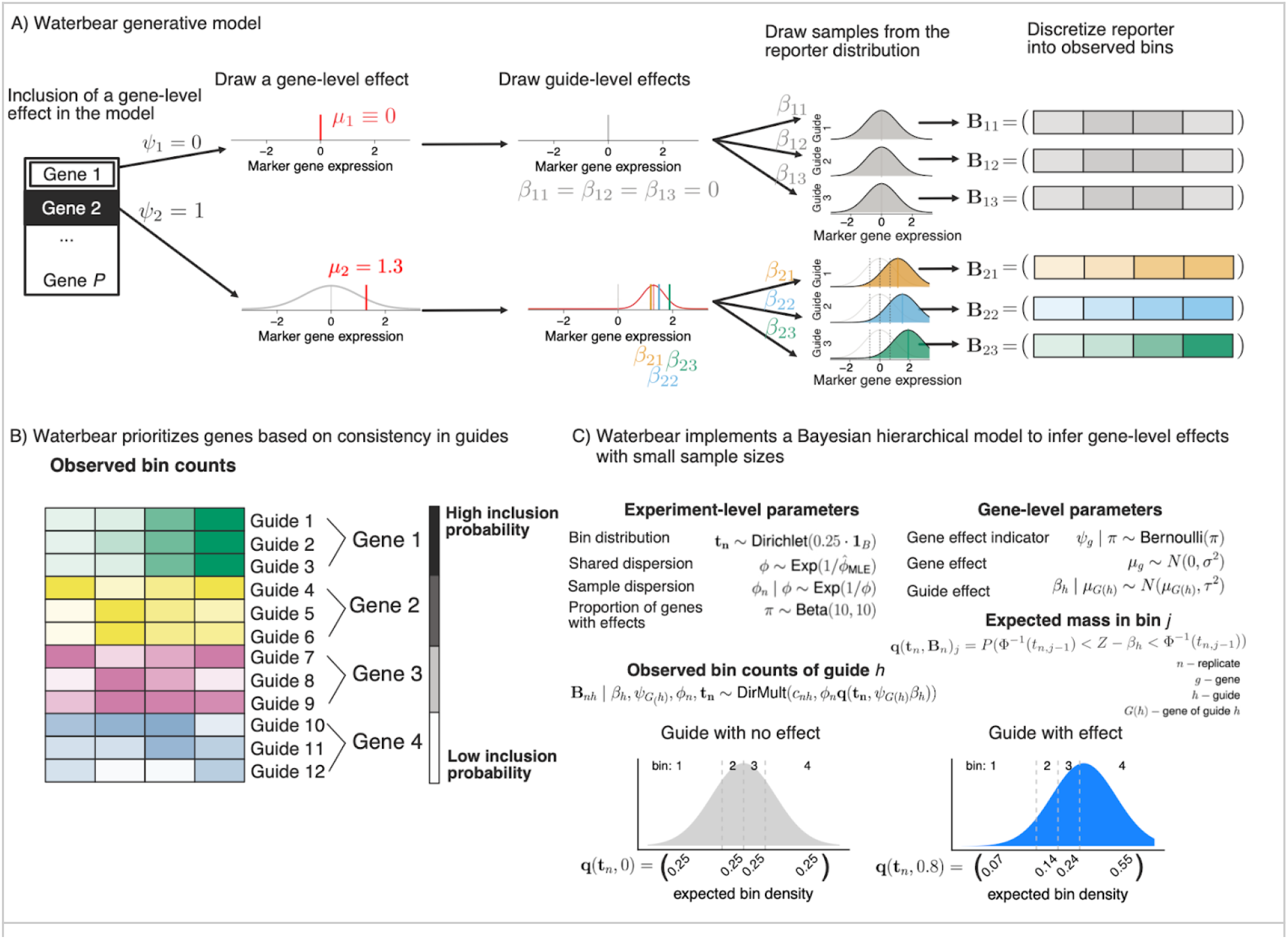
A statistical model for CRISPR FACS screen data. A) Overview of the generative model. First, the gene level perturbation effect is assessed as either having a no effect (top row) or having an effect (bottom row) on the marker gene and the effect size established as *μ*1. For genes with an effect on the marker, the individual gRNA effect sizes are chosen centered around this gene effect. Each gRNA is assumed to shift the expression of the marker gene to center on the gRNA’s effect size. Finally, each marker is discretized into four FACS bins with the counts in each bin reflecting the amount that the marker distribution was shifted. B) Multiple gRNAs that have consistent effects on a given gene increase the probability that perturbation of that gene affects the level of the marker. In this example, gRNAs 1-3 target Gene 1 and show a consistent up regulation of the marker, thus, Gene 1 has the highest posterior inclusion probability. gRNAs 4-6 have more noise and less overall up regulation, so Gene 2 shows a slightly lower posterior inclusion probability than Gene 1. C) Waterbear implements a Bayesian hierarchical model to infer gene-level effects with small sample sizes by sharing parameters within genes and across the experiment. Additionally, Waterbear improves inference by modeling the unobserved FACS distribution (bottom row). When there is no shift, the guide distribution is modeled to look like a control guide (bottom left). When there is an effect, the bin proportion and thus bin counts are shifted (bottom right).

When a gene has no effect (*ψ_i_* = 0), all of the gRNAs targeting this gene should be similar to the null marker distribution with effects being shrunk towards zero. Thus, deviations in the observed sequencing counts between bins represent noise in the experiment. In particular, control gRNAs enable us to force *ψ_i_* = 0 which enables a direct estimation of the experimental noise.

When a gene has an effect ( *ψ_i_* = 1) the model allows the guide level effect estimates to vary and we employ a hierarchical process which assumes gRNAs targeting that gene behave similarly. The true gene-level effect is drawn from a Gaussian effect distribution enabling a large range of true effects, as is common in Bayesian models due to its flexibility and modeling convenience ^17^. Then, each gRNA has its own random effect centered around the gene-level effect distribution. Each gRNA thus results in a different marker distribution that is shifted around the gene-level effect, resulting in a gRNA-specific gRNA-marker bin count distribution.

As a result of this two-class approach, rather than focusing on gRNA- or gene-level p-values, Waterbear prioritizes genes where the individual gRNAs targeting that gene produce consistent shifts in the marker distribution (Figure 2B). In particular, a gene is considered a notable target if the inferred posterior inclusion probability (PIP), Pr[*ψ_i_* = 1 | data] , is sufficiently close to one. Genes with high PIP have gRNAs with consistent, non-zero effects and genes with low PIP have gRNAs with effects close to zero. As the count of inconsistent gRNAs targeting a gene rises, the PIP will decrease, indicating higher uncertainty in the relationship between the target gene and the marker. Statistically, this is one of the major contributions of Waterbear; rather than aggregating results from each guide, Waterber fits a holistic model in which the data help infer whether the “effect” class or the “no effect” class is more consistent over all guides targeting a gene.

Importantly, Waterbear learns experimental parameters including the sizes of the sorted FACS bins using either the counts of the control gRNAs or, in the absence of controls, by assuming that a subset of gRNAs do not have an effect (Figure 2C). Experimentally, FACS bin sizes can shift during long sorts and opposing bins (e.g. bin one and bin four) might not always be collected in equal proportions. Using the control gRNA counts in each bin, Waterbear infers for each replicate what bin cutoffs divide the marker distribution to produce the observed count distribution.

### Waterbear has relatively high sensitivity while controlling the false discovery rate

To establish the performance of Waterbear and compare it to other methods, we used the cell-level view of the generative model which includes a number of tunable parameters (Supplementary Section 2). We performed simulations and analyzed the results with the collapsed “gene-level” view of Waterbear, MAGeCK, and MAUDE (Methods) ^12,13^ to observe how changing these parameters affect each method’s ability to detect hits.

To assess each method’s calibration, we compared the estimated false discovery rate (FDR) to the true FDR. Ideally, the true FDR should be at or below the estimated FDR. We first ran simulations across a range of coverage levels with 10% of the gRNAs in the library having a true effect on the marker. At all tested coverage levels, Waterbear and MAGeCK maintained lower true FDRs at the estimated 10% FDR (Figure 3A). In contrast, MAUDE’s true FDR was nearly 50% across all coverage levels. We also evaluated the calibration with low cell:gRNA ratios while varying the MOI. Increasing the MOI resulted in a higher effective coverage without having to increase cell numbers (Figure 1B). Again, MAGeCK and Waterbear controlled the FDR, while MAUDE had a highly inflated FDR for nearly all MOI levels (Figure 3B).

**Figure 3:**
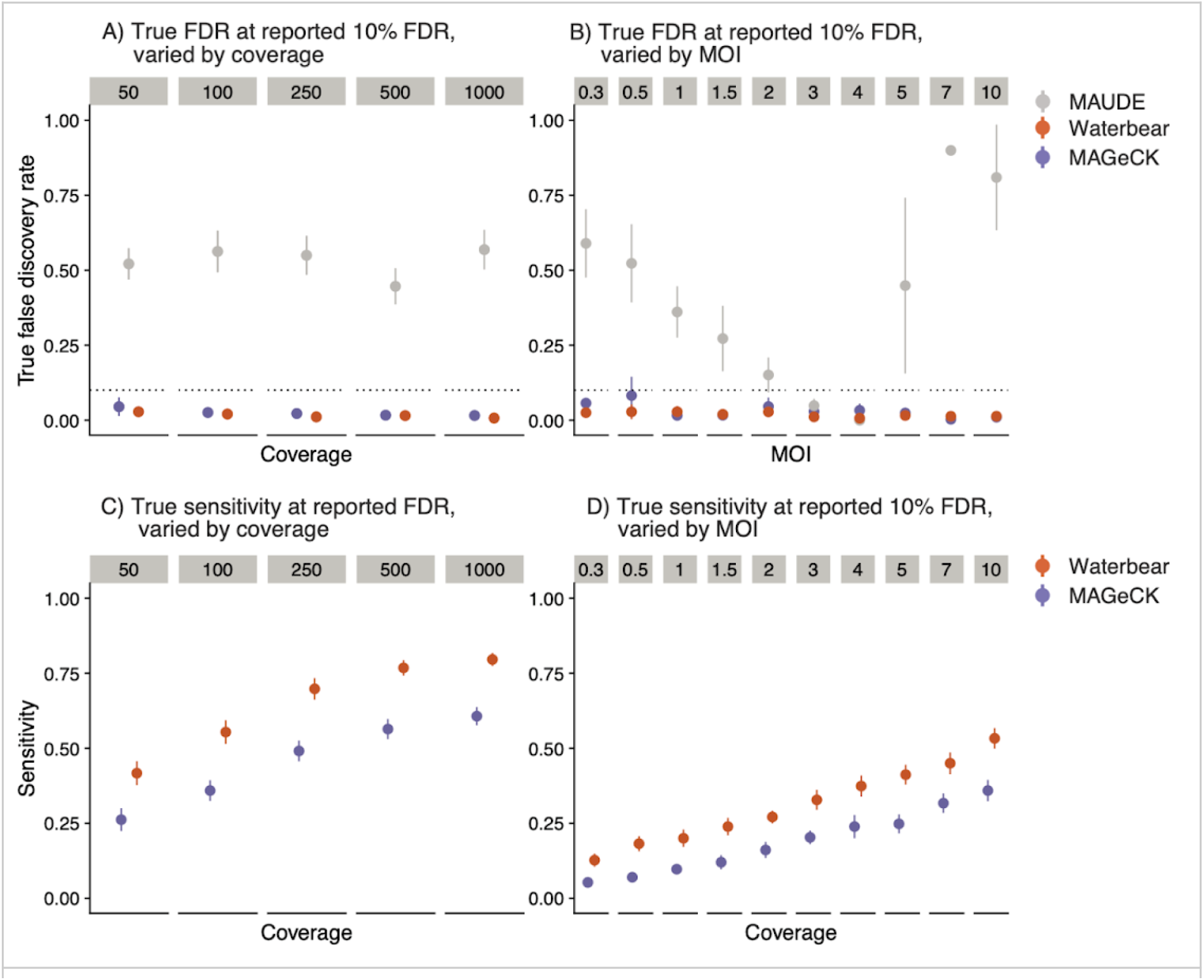
Waterbear has relatively high sensitivity while controlling the false discovery rate. Sensitivity and calibration plots comparing Waterbear, MAGeCK, and MAUDE on simulated screen data. For each method, we consider a hit to be ‘significant’ if the estimated FDR is less than q = 0.10. Since Waterbear produces posterior inclusion probabilities, we consider a test to be significant if PIP > 1 - q and the gene effect size (1 - q) credibility interval does not include zero. True FDR of the methods across various coverage levels (A) and various MOIs (B). Sensitivity of the methods across various coverage levels (C) and various MOIs (D).

Experiments are often collected under less ideal conditions than simulations. Given that Waterbear learns most parameters from the observed data rather than making assumptions about the experiment, we thought that it should be more robust than existing tools under non-ideal conditions. For instance, since MAGeCK was not designed to analyze FACS-based screens, using it for this type of analysis requires the implicit assumption that all FACS bins are equally sized with each other and between donors. While MAUDE supports uneven bin sizes, it requires the user to manually specify the exact bin sizes collected. In contrast, for each sample Waterbear learns the sizes of individual FACS bins directly from the data. Therefore, we expected that Waterbear would outperform both MAGeCK and MAUDE in situations where the bin sizes were misspecified. In simulations with uneven bin sizes, Waterbear and, surprisingly, MAGeCK controlled the FDR, while MAUDE did not (Supplementary Figure 2.8). It is important to note that since MAUDE does not estimate the bin sizes, we input the true simulated bin sizes for MAUDE, while letting Waterbear estimate the bin sizes and MAGeCK normalize the counts. These results show that MAGeCK’s RNA-seq style normalization still performs well in this setting.

We next tested which methods had the highest sensitivity while controlling the FDR. Ideally, sensitivity should be close to one, while the FDR is close to zero. Consistently across different coverage levels, MOI, and bin sizes, Waterbear had the highest sensitivity while controlling the FDR (Figures 3C-D). While MAUDE technically had the highest sensitivity, this came at a high false discovery cost, as nearly half of the calls were false positives. Despite not being designed for FACS screens, MaGeCK had low FDR and still had relatively high sensitivity, albeit, nearly always second to Waterbear.

### Waterbear simulations suggest high sensitivity is maintained at low cell counts and high MOI

As discussed previously, we posited that the gRNA coverage, FACS bin configuration, and MOI have the largest impacts on experiments, so we focused on these parameters in our simulations. Given that we know the ground truth for these simulations, we can calculate how changing any of these parameters affects the sensitivity of the screen. Our cell-level generative model and effect size distributions are detailed in Supplementary Section 2.

Increasing the gRNA coverage with fixed MOI enables a more accurate quantification of how perturbations affect the marker (Figure 4A). However, at high coverage levels, a large increase in cell number yields only diminishing returns for quantifying gRNAs. This result suggests that identifying the minimum coverage needed to detect significant hits would have a meaningful impact on reducing the resources required for FACS screens. For example, Figure 4A shows that 38% of true hits will still be detected with only 50X coverage at a true FDR of 0.05.

**Figure 4:**
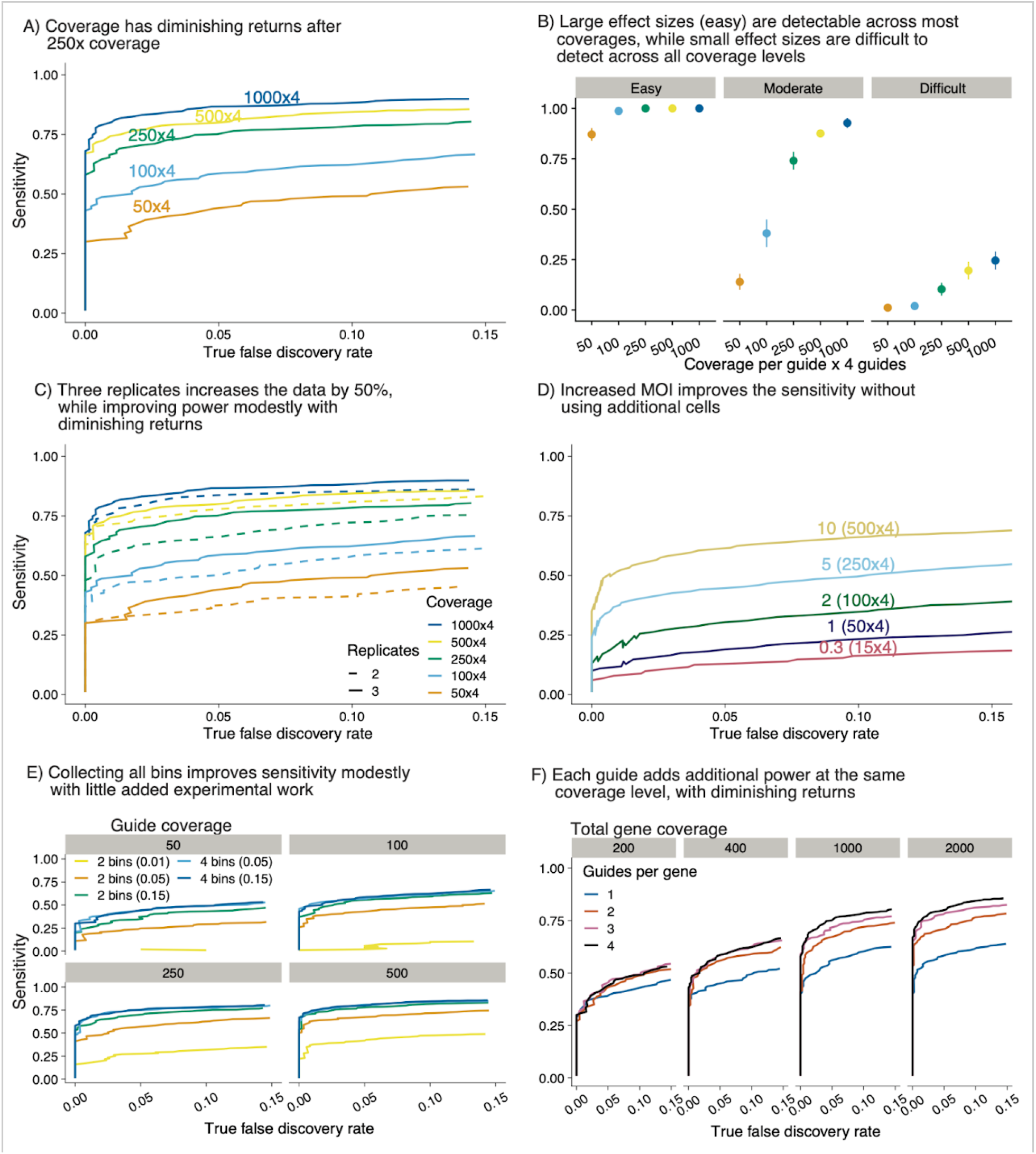
Waterbear simulations suggest high sensitivity is maintained at low cell counts and high MOI. A) As gRNA coverage increases, sensitivity approaches saturation. In this simulation there are 4 gRNAs per gene, 1,000 genes, and 10% of all genes have an effect. Each line indicates the coverage at the gRNA level. B) Sensitivity of (A) broken down by the size of effect that the gRNA perturbation has on the marker expression levels. C) Comparison of 2 replicates versus 3 under the same conditions as (A). D) Effect of different MOIs on sensitivity. In this example we simulated 50,000 cells. E) Waterbear can be used with data from all 4 bins or by imputing unobserved middle bins (2 bin mode). Sensitivity assessed at 4 coverage levels. F) Effective coverage at the gene level stratified by the number of gRNAs. Each line indicates a different gRNA configuration and each subpanel indicates a fixed total gene level coverage.

Furthermore, in this scenario, almost 80% of large effect size hits, defined as having an effect size greater than 0.2 standard deviations, will be detected (Figure 4B). Therefore, the strongest hits in a given screen will be detected even when the coverage is as low as 50X per gRNA. Increasing the coverage increases the number of hits detected and allows the detection of hits with a smaller effect size. However, the sensitivity starts to saturate around 1000X coverage, suggesting that it is not worthwhile to collect additional cells.

We next asked, given limited experimental resources, would it be better to collect a third biological replicate or to collect two replicates with higher coverage. At lower coverage levels, adding a third replicate only increased the mean sensitivity roughly half as much as doubling the coverage of the existing two replicates (Figure 4C). At higher coverage levels, the two replicates already start to saturate the mean sensitivity, suggesting that a third replicate would only have minimal benefit. To confirm this result on real data, we down-sampled data from Figure 5 to two replicates and found comparable sensitivity across the down-sampled counterparts and that the top hits in every configuration replicated (Supplementary Section 3.1).

**Figure 5:**
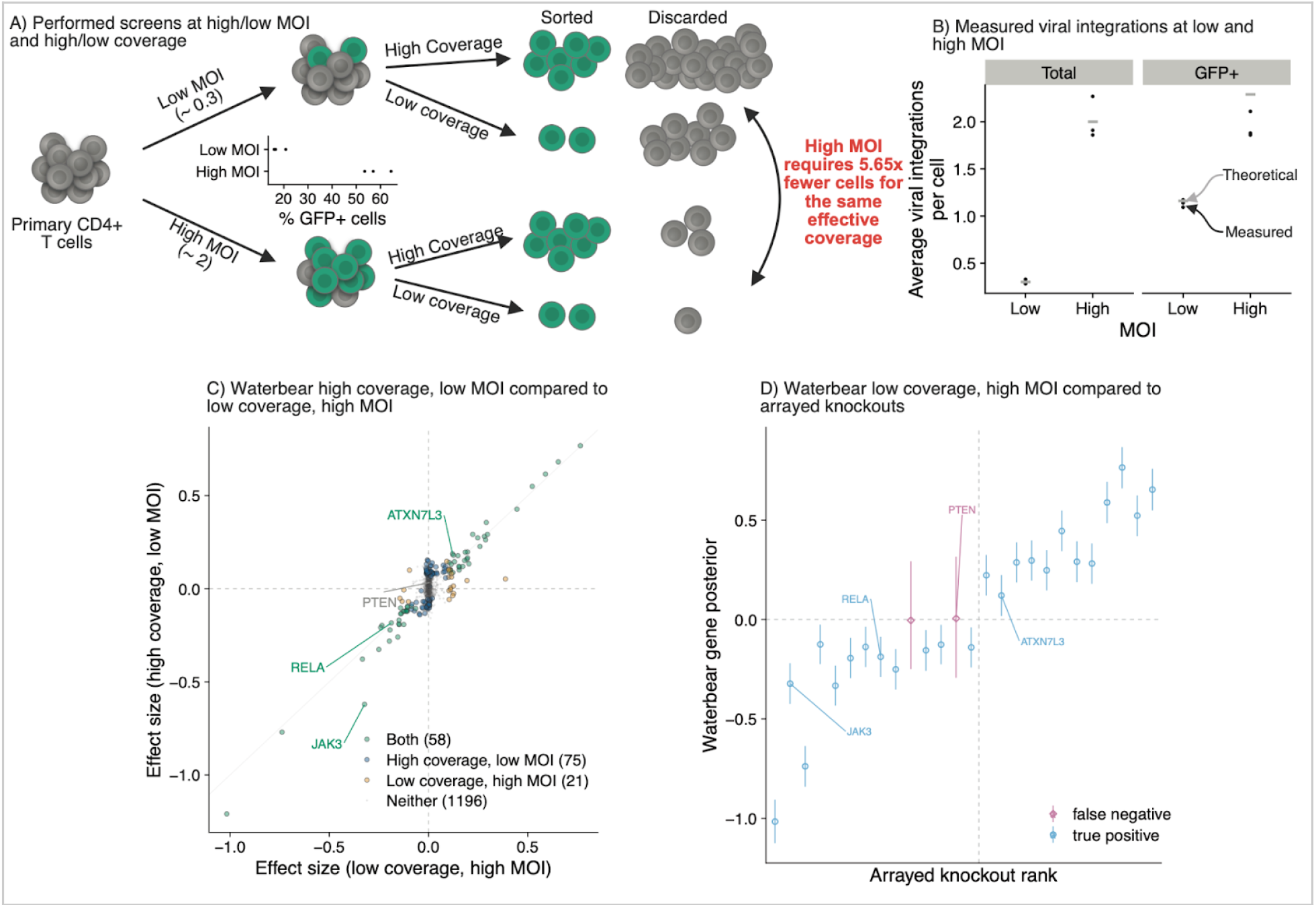
Experiments validate that high sensitivity is maintained at low cell counts and high MOI. A) CRISPR FACS screens to identify regulators of IL2RA were performed at low (∼0.3) and high (∼2) MOI and low and high coverage. Low coverage (average of 195x coverage for MOI 0.3, 208x coverage for MOI 2) and high coverage (average of 1662x coverage for MOI 0.3, 1180x coverage for MOI 2) n = 3 donors. The relative number of cells collected for each condition is shown. B) Quantification of the number of viral integrations per cell using droplet digital PCR in the unsorted cells (total) or after sorting cells that express GFP in the gRNA lentiviral construct (GFP+). C) Comparison of screens hits from the high coverage, low MOI screen vs low coverage, high MOI screen analyzed using Waterbear. D) Experimentally validated regulators of IL2RA were detected as screen hits in the low coverage, high MOI screen using Waterbear.

Commonly, screens are often performed using a low MOI of 0.3 - 0.5 to minimize the number of cells that contain more than one gRNA. However, with an MOI of 0.3, approximately 74% of cells will receive no gRNAs during the viral infection, which often limits the number of gRNAs that can be screened, especially when dealing with a limited number of starting cells. Therefore, at a fixed coverage for the experiment, even modestly increasing the MOI drastically reduces the number of cells needed (Figures 1Bii). We fixed the number of cells at 50,000 with a 6,000 gRNA library. Increasing the MOI served to effectively increase the coverage, while not having any negative effect on mean sensitivity even as the MOI approached 10 (Figure 4D). This trend is also observed when going to higher numbers of total guides and different proportions of guides with an effect (Supplementary Section 2.2). To check if the number of controls affected the inference, we also performed many of the simulations with 10, 100, and 1000 controls (Supplementary Section 3.2) which verified that this does not affect the final result. Thus, increasing the MOI, even a marginal amount, can greatly improve sensitivity.

While increasing the MOI will result in some cells containing random combinations of multiple gRNAs, in our previous screens less than 10% of the gRNAs had a statistically significant effect. In similar screens, the majority of cells will contain combinations of gRNAs where none or only one of the gRNAs have an effect (Supplementary Section 4). We define cells containing “gRNA collisions” as cells which contain two or more gRNAs that exhibit an effect on the marker distribution, but even at moderate MOIs gRNA collisions are rare. If a gRNA has an effect then cells containing both this gRNA and random mostly no-effect background gRNAs should still end up enriched in either the low or high FACS bins. However, if a gRNA has no effect and it is randomly paired with mostly no-effect background gRNAs, cells containing these no-effect gRNAs will end up equally distributed between the high and low FACS bins.

Many FACS screens only collect the outer two bins. However, we expected that sequencing four bins spanning the full distribution would increase the gRNA coverage data that Waterbear uses for inference without requiring additional input cells. We asked, given the same number of cells: How moving from two bins to four bins affects the sensitivity to detect hits (Figure 4E)? With 15% outer bins, four bins is almost always preferable, however, the increase in sensitivity is relatively minor. However, performance greatly degrades if one only collects two outerbins at the tail of the extremes of the marker distribution (5% and 1% outer bins), but is maintained if one also collects the inner additional bins. In summary, sequencing four bins provides consistent results across coverage levels tested and is preferable when coverage is low (<= 50X gRNA coverage).

While these simulations suggest that the cumulative coverage is an important driver of sensitivity when four gRNAs target the same gene, it is unclear whether four gRNAs are necessary. We next explored how the number of gRNAs affects the sensitivity when the coverage is fixed per gene, under the assumption of high-quality gRNAs. To test the effect of the number of gRNAs targeting a gene, we performed an analysis where the total gene coverage was fixed, but was achieved using 1, 2, 3, or 4 gRNAs (Figure 4F). For example, at 1000X coverage, 1 gRNA represents 1000 cells for that gRNA, whereas in the 4 gRNA case, it represents 250 cells per gRNA. Using one gRNA almost always has less power than other configurations. As coverage increases, more gRNAs are helpful up to about three guides. This result is likely due to the additional evidence against the null as each additional gRNA can be thought of as its own “experiment” during inference, akin to a meta-analysis.

While these simulations suggest that increasing the coverage by many different means will increase the ability to detect hits, they also provide a path to reducing the number of cells and thus enabling novel screens in systems previously limited by the number of cells available. In particular, increasing the MOI while capturing four bins can increase the overall coverage with relatively small experimental burden and no additional cells.

### Experiments validate that high sensitivity is maintained at low coverage and high MOI

We previously performed CRISPR FACS-based screens in primary human T cells to identify the upstream regulators of *IL2RA*, an important cell surface receptor implicated in numerous autoimmune diseases ^3,18,19^. However, these screens were expensive and experimentally demanding because they were performed at 640-2273x coverage and each screen required 100-290 million primary cells. However, our simulations suggest that we could have identified the top hits using much lower coverage and a higher MOI. To confirm these results, we repeated the screen using an MOI of 0.3 (low MOI) and 2 (high MOI). For both MOI conditions, we collected the cells from 3 donors across 4 FACS bins at high coverage (average of 1662x coverage for MOI 0.3, 1180x coverage for MOI 2) and at low coverage (average of 195x coverage for MOI 0.3, 208x coverage for MOI 2) (Figure 5A).

To confirm that the cells were infected at the desired MOI, we quantified the genomic integration copy number for the gRNA lentiviral construct. We used droplet digital PCR to quantify the number of copies of GFP, which is part of the gRNA lentiviral construct, relative to the number of copies of the control gene RPP30 in the cells. The average copy number of GFP in the population ranged from 0.28 to 0.33 for the MOI 0.3 condition and from 1.9 to 2.3 for the MOI 2 condition across the 3 donors, closely matching the theoretical copy number for each condition (Figure 5B). These data suggest that the cells were infected at the desired MOIs and that the majority of cells in the high MOI condition contained more than one gRNA per cell. Given that the majority of cells in the low MOI condition contained no gRNAs, one would need ∼5.5 fold more cells to obtain the same effective coverage as the high MOI condition.

We next compared the significant hits in the high coverage, low MOI condition compared to the low coverage, high MOI condition using Waterbear. Despite having multiple gRNAs per cell and being collected at 4.6-fold lower coverage, the top hits were highly correlated between the two conditions (Figure 5C). We previously validated 26/33 hits from our original screen by performing individual knockouts and directly measuring the effect on IL2RA protein levels using flow cytometry ^3^. We used these 26 genes as a set of high confidence positive controls (Supplementary Section 3.2). Waterbear detected 24 out of 26 of these validated hits in the low coverage, high MOI screen (Figure 5D). Together, these results experimentally validate both the predictions regarding coverage and MOI from our simulations with Waterbear as well as demonstrate that Waterbear is a powerful tool to analyze these screens, even under challenging conditions.

Given that Waterbear outperformed MAGeCK and MAUDE in our simulations, we wanted to compare Waterbear’s sensitivity to these tools under real conditions. On the low coverage, high MOI screen MAGeCK detected 17 out of 26 of these validated hits (Supplementary Figure 3.6). MaGeCK’s lower sensitivity to Waterbear’s (24/26) is consistent with observation that in general MaGeCK generally calls fewer hits with lower sensitivity; Waterbear called 79/1350 and MAGeCK called 31/1350 genes significant. MAUDE reports many more signals (406/1350 and detects 25/26 hits, Supplementary Figure 3.7), however we regard these as inflated given the simulations showing poor calibration at 10% FDR. These results further validate that Waterbear maintains high sensitivity, without seemingly over-calling relationships.

## Discussion

Genetic screens are a powerful approach to link genes to phenotypes. Coupling CRISPR perturbations with FACS in mammalian cells enables mapping the genetic basis of many biological processes and regulatory relationships. Early CRISPR screens were performed in abundant cell lines ^20–22^, but increasingly, these screens are being performed in rare primary cell types or *in vivo* models with limited cell numbers ^4,9,23,24^. In these screens, researchers must often make choices about experimental conditions based on their intuition as there have not been good guidelines to inform how changing experimental parameters affect the ability to identify hits. Furthermore, changes such as lowering gRNA coverage or increasing the MOI increase the statistical challenge of identifying hits.

We solve these experimental design and analysis gaps by introducing Waterbear, an end-to-end experimental design and inference procedure for CRISPR FACS screens. Our cell-level generative implementation enables exploration of experimental parameters such as effect size distribution, gRNA distribution, and MOI. In conjunction with a “gene-level” version of this model for inference, Waterbear enabled us to show that (1) sensitivity saturates relatively quickly, and thus if using four bins can be dropped from about 1,000X coverage to 250X coverage with little loss in sensitivity, (2) increasing the MOI modestly from 0.3 to 2 improves effective coverage while greatly reducing the number of input cells needed, (3) the number of gRNAs targeting each gene can be reduced from four to three while achieving similar sensitivity.

Overall, our results demonstrate that the number of cells for such screens can be reduced, enabling one to assay significantly smaller cell populations. The prevailing view has been that such screens should be performed with only a single perturbation per cell. These guidelines likely emerged from siRNA screens where there are many more off-target effects. Consistent with other recent reports ^25–27^, we demonstrate that increasing the MOI greatly reduces the number of uninfected input cells that are thrown away during screening, while having a similar sensitivity as the low MOI screen. Coupled with our results that coverage and gRNA number can be reduced without impacting sensitivity, these guidelines should reduce the resources and effort required to perform CRISPR FACS screens and enable a next generation of CRISPR screens in rare cell types and with *in vivo* models that will be essential to understand many disease-relevant processes.

Equipped with knowledge of the important experimental aspects, we developed a hierarchical statistical model that enables principled inference of guide-effects with replicates. The unique approach models the unobserved FACS marker and connects it to the observed sequencing counts. This key hierarchy is what sets the Waterbear method apart from existing methods and makes it adaptable for other FACS-based sequencing experiments. The statistical challenge lies in how one connects the unobserved marker distribution to the FACS bins, while maintaining the correlation structure in the bins and not treating each bin independently, but rather, as samples joint from an unobserved marker. Concretely, this is modeled through our function *q*(·) in the methods section.

Importantly, the Waterbear model infers experimental parameters and is robust in small sample settings which are common in these screens. We do so by “shrinking” estimates in the hierarchy, thus jointly modeling experimental parameters across replicates or within replicates when appropriate. To address the concerns of model assumptions and general performance, we additionally demonstrated that our model outperforms tools commonly used for these types of analyses (MAGeCK and MAUDE) in many different experimental settings and on real biological data.

Through both simulations and follow-up experiments, we have demonstrated an approach of model driven experimental design followed by experimentally guided inference models that can be seen as a vignette for other similar screens. For example, both the experimental design and inference framework can be modified to inform design and inference of proliferation screens, *in vivo* screens, scRNA-seq screens, and potentially multi-ome readout screens.

## Data Availability

The raw sequencing files generated during this study are available at GEO: GSE242880.

## Code Availability

The code to reproduce all of the analyses in this paper can be found at: https://github.com/pimentel/waterbear_analysis. The pipeline tool Snakemake ^28^ was used to run Waterbear, MAGeCK, and MAUDE.

## Methods

### Sample collection

This study was approved by the University of California, San Francisco (UCSF) Committee on Human Research and Stanford University Panel on Medical Human Subjects (IRB#53302) and written consent was obtained from all donors. Primary human T cells were obtained through consented Leukopaks (STEMCELL) (Catalog #70500.2).

### Isolation, culture and expansion of human CD4+CD25-effector T cells

PBMCs from Leukopaks (STEMCELL) were diluted 1:1 with PBS containing 2% FCS and 1mM EDTA and spun at 500g for 10 minutes. The StemCell EasySep Human Isolation Kit (Catalog # 18063) was used to isolate CD4+CD25-effector T cells from washed PBMCs while excluding CD4+CD25+ regulatory T cells. Isolated cells were then stimulated with Immunocult Human CD3/CD28/CD2 T Cell Activator (STEMCELL, Cat #10970) at 6.25 uL per 1E6 cells and grown in RPMI with 50 U/mL IL-2 (Amerisource Bergen, Cat #10101641) at a concentration of 1E6 cells/mL.

### Pooled CRISPR screens

Pooled CRISPR screens were performed as in Freimer et al. ^3^.

#### Lentiviral transduction

Approximately twenty-four hours post stimulation, lentivirus containing the sgRNA library was added directly to cultured T cells at various multiplicity of infections (MOIs). After an additional twenty-four hours, the media was changed.

#### Cas9-ribonucleotide protein (RNP) preparation

Cas9 (MacroLab, Berkeley, 40 µM stock) ribonucleoprotein complex was delivered into the cells using a modified Guide Swap technique ^29^. Lyophilized Dharmacon Edit-R crRNA Non-targeting Control #3 (Dharmacon, Cat #U-007503-01-05) and Dharmacon Edit-R CRISPR-Cas9 Synthetic tracrRNA (Dharmacon, Cat #U-002005-20) were resuspended at a stock concentration of 160 uM in 10 mM Tris-HCl (pH 7.4) with 150 mM KCl. They were mixed at a 1:1 ratio and incubated at 37°C for 30 minutes. A single-stranded donor oligonucleotide (ssODN; sequence: TTAGCTCTGTTTACGTCCCAGCGGGCATGAGAGTAACAAGAGGGTGTGGTAATATTACGGTACCGAGCACTATCGATACAATATGTGTCATACGGACACG) was then added at a 1:1 molar ratio of the final Cas9-Guide complex and the solution was mixed well by pipetting. The solution was incubated for an additional 5 minutes at 37°C. Cas9 protein was then added slowly at a 1:1 volume and incubated at 37°C for 15 minutes.

#### Electroporation

Approximately twenty-four hours after viral transduction the cells were centrifuged at 100*g* for 10 minutes and then resuspended in room temperature Lonza P3 electroporation buffer (Lonza, Cat #V4XP-3032) at 1-2E6 cells per 17.8 µL. For every 17.8 µL of cells, 7.2 µL of the RNP-ssODN complex was added and the solution was mixed well. 23 uL of the cells-RNP-ssODN mixture was added to each well of a 96 well electroporation cuvette plate (Lonza, Cat #VVPA-1002), and nucleofected using the pulse code EH-115. Immediately after electroporation, 90 µL of warm media was added to each well and incubated at 37°C for 15 minutes. Cells were then pooled and grown at a concentration of 1E6 cells/mL.

### Screen phenotyping and cell sorting

Cells were collected for analysis 6 days after electroporation. Cells were stained for IL2RA using an APC fluorescent antibody (Tonbo, Cat #20-0259-T100) at a 1:25 dilution according to the manufacturer’s protocol. GFP positive cells were sorted into 4 bins based on IL2RA protein levels using a BD FACS Aria II and FACSDiva version 8.0.1. Bulk GFP positive and negative population were also collected for ddPCR assessment.

### GFP copy number assessment using Droplet Digital PCR

Following genomic DNA purification, an aliquot was reserved for ddPCR analysis. For each sample, 10 ng of purified genomic DNA was added to a reaction consisting of 10 µL of ddPCR Supermix for Probes (Bio-Rad, Cat #1863024), 1 µL of MseI (New England Biolabs, Cat #R0525S), 1 µL of both the reference and target primer assays, and water to a total volume of 20 µL. A Bio-Rad validated HEX ddPCR Copy Number Assay targeting the reference gene, RPP30, (Cat #10031243) was used in addition to a custom FAM Copy Number Assay targeting GFP (Bio-Rad, Cat #10042958). After assembling the ddPCR reactions, the samples were emulsified in oil droplets using the QX200 Droplet Generator (Bio-Rad, Cat #1864002) following the manufacturer’s instructions. In brief, a DG8tm Cartridge (Cat #1864008) was inserted into a DG8 Cartridge Holder (Cat #1863051) and each 20 µL reaction mixture was transferred to a well, followed by 70 µL of Droplet Generation Oil for Probes (Cat #1863005). The cartridge was covered with a gasket (Cat #1863009) and loaded into the Droplet Generator. Following emulsification, the droplets were transferred to a 96 well plate (Cat #12001925). This process was repeated until all samples were transferred to the plate, which was then sealed using a pierceable heat seal (Cat #1814040) and the PX1 PCR Plate Sealer (Cat #1814000). DNA was was fragmented and amplified using the BioRad C1000 Thermocycler (Cat #1851196) programmed to the following specifications: 95°C for 10 minutes, followed by 40 cycles of 94°C for 20 seconds and 57°C for 1 minute, and ending with 98°C for 10 minutes and a final 4°C hold until data acquisition. Each step was performed with a ramp rate of 2°C/sec until the final cool down at 1°C/sec. Amplification was determined with the QX200 Droplet Reader (Cat #1864003) using the Copy Number Variant (CNV) assay on the QuantaSoft™Software. Positive and negative populations in each channel were manually defined using the oval tool.

### Lentiviral production

14E6 HEK 293T cells were seeded in a 15 cm tissue culture dish (Corning, Cat #430599) in Opti-MEM (UCSF CCF, Cat #CCFAC008) approximately twenty-four hours prior to transfection. Cells were transfected with the sgRNA library plasmid, and two lentiviral packaging plasmids, pMD2.G (Addgene, Cat #12259) and psPAX2 (Addgene, Cat #12260) using Lipofectamine 3000 (Lifetech, Cat #L3000075). Cells were incubated for 5 hours at 37°C. The media was then replaced with fresh Opti-MEM containing ViralBoost at 1x (Alstem, Cat #VB100). The cells were cultured for approximately twenty-four hours and then the media was collected and spun down at 300*g* for 5 minutes to remove cellular debris. The media was then filtered using 0.45-µm filter and one volume of cold Lentivirus Precipitation Solution (Alstem, Cat #VC125) was added for every four volumes of lentivirus-containing media. Samples were mixed and then put at 4°C overnight. The viral media was then spun in a centrifuge at 1500*g* for 30 minutes at 4°C, followed by a second spin at 1500*g* for 5 minutes to concentrate the virus. The viral pellet was then resuspended in 4°C PBS (Fisher Scientific, Cat #10010049) at a 1:100 dilution of the original media volume. The concentrated virus was stored at -80°C until use.

### Culture media

Cells were grown in RPMI (Sigma, Cat # R0883) with 10% FCS (Sigma, Cat # F0926), with 100U/mL Pen-Strep (Gibco, Cat # 15140-122), 2mM L-Glutamine (Sigma, Cat # G7513), 10mM HEPES (Sigma, Cat # H0887), 1X MEM Non-essential Amino Acids (Gibco, Cat # 11140-050), 1mM Sodium Pyruvate (Gibco, Cat # 11360-070), and 50 U/mL IL-2 (Amerisource Bergen, Cat #10101641) at a concentration of 1E6 cells/mL.

### Genomic DNA extraction and preparation for next generation sequencing

After sorting, cells were washed with PBS, counted, and resuspended at up to 5E6 cells per 400 µl of lysis buffer (1% SDS, 50 mM Tris, pH 8, 10 mM EDTA). 16µl of NaCl (5M) was added and the sample was incubated overnight at 66°C. Then 8µl of RNAse A (10mg/ml) (Thermo Scientific, Cat #EN0531) was added and incubated at 37°C for 1 hour. Then 8µl of Proteinase K (20mg/ml) (Thermo Scientific, Cat #AM2548) was added and incubated at 55°C for 1 hour. A phase lock tube (Quantabio, Cat #2302820) was prepared for each sample and then 400µl of Phenol:Chloroform:Isoamyl Alcohol (25:24:1) was added to each tube. 400µl of the sample was then added and the tube was shaken vigorously. The sample was centrifuged for 5 min at max speed at room temperature. The aqueous phase was transferred to a low-bind Eppendorf tube (Eppendorf, Cat #022431021). 40µl of Sodium Acetate (3M), 1µl GlycoBlue (Invitrogen, Cat # AM9515), and 600µl of room temperature isopropanol were added to the sample. The sample was stored at -80°C for 30 minutes and then centrifuged for 30 minutes at max speed at 4°C. The pellet was washed with fresh 70% room temperature Ethanol and allowed to air dry for 15 minutes. Pellets were then resuspended in Zymo DNA elution buffer (Zymo, Cat No: D3004-4-10), and then incubated at 65°C for 1 hour to completely dissolve the genomic DNA.

sgRNAs were amplified from the genomic DNA as initially described by Joung et al. ^30^. Up to 2.5 µg of genomic DNA was added to each 50 µL PCR reaction with 25 µL of NEBNext Ultra II Q5 master mix (NEB, Cat #M0544L), 1.25 µL of the 10 µM forward primer and 1.25 µL of the 10 µM reverse primer, and H2O to 50 uL. The reaction was then amplified with the following cycling conditions: 98°C for 3 minutes, followed by 23 cycles at 98°C for 10 seconds, 63°C for 10 seconds, and 72°C for 25 seconds, and finally 2 minutes at 72°C. The amplicons were cleaned with Sera-Mag Speed Beads (Cytiva, Cat #65152105050250) used at a 1X v/v ratio. The concentration of each sample was then measured using the Qubit dsDNA high sensitivity assay kit (Thermo Fisher Scientific, Cat #Q32854) and the successful removal of adapter dimers confirmed with the 4200 TapeStation system (Agilent, Cat# G2991BA). Samples were then sequenced on an Illumina HiSeq 4000 (Illumina, Cat #15017872) using a custom sequencing primer.

### Statistics and analysis

#### A cell-level generative model for FACS screens

For Figures 3 and 4, we simulate from a cell-level generative model. The full description of the cell-level generative model is in Supplementary Section 1.1, but we briefly describe the cell-level process here. First, a number of guides is chosen according to a Poisson distribution (MOI).

Next, the guides integrating with this cell are chosen at random, according to a Dirichlet. If more than one of the included guides has an effect, we assume additive, linear effects, and no interaction effects. A value from the resulting marker distribution is drawn and the corresponding guide bin count is incremented. The final observation is a Dirichlet Multinomial centered around the total bin counts corresponding to a noisy observation of the entire process.

#### Parameters for simulations

The guide representation proportion is modeled as 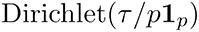 where *τ* > 0 represents the dispersion and *p* represents the number of guides. The parameter *τ* was learned from the GFP+ population on real data. Further, effect sizes were also learned on this data set. See Supplementary Section 2 for full details.

#### A model for gene-level inference

Waterbear implements a hierarchical Bayesian model to infer the latent effect sizes at the gene-level. In order to allow inference with small sample sizes that are common in screens, we use various shared priors in the hierarchy. We begin with the observed bin counts in sample *n*, resulting from guide *h* by,

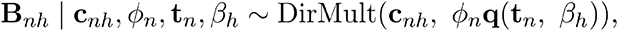

where c*_nh_* is the guide coverage, *ϕ_n_* is the sample specific dispersion, and q(*t_n_*, *β_h_*) takes the null probability mass in each bin (t*_n_*) and the guide effect size (*β_h_* ) and returns the probability mass function across the bins. In particular, t*_nj_* is the mass in bin *j* under the null marker and thus we model the joint over t*_n_* as a Dirichlet. If we assume the marker distribution to be Normal(*β_h_*, 1), the mass in bin *j* is defined as,

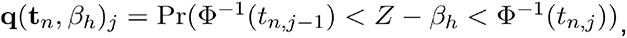

where we define t*_n,_*_0_ = 0 and t*_n,_*_5_ = 1 (under a 4-bin experiment), *Z* is a standard normal random variable, and 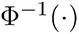 is the inverse CDF of a standard normal.

To draw guide level effect sizes, first we must decide if the gene is included in the model *ψ_g_* =1 or if it is not (*ψ_g_* =0). Guide-level effect sizes are drawn from a prior centered at the latent gene effect, *μ_g_* if the gene is included in the model or a point mass at 0 if it is not according to a spike-and-slab prior,

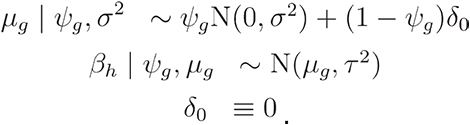

At the population level, the parameter *π* dictates the proportion of genes that are allowed to be included in the model (*ψ_g_*). Thus, 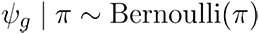. Of note, this prior encourages guide-level effects if there is evidence with many guides. A full specification of all priors and hyper parameters can be found in Supplementary Section 1.2. Posterior sampling is performed with a MCMC sampler implemented in NIMBLE ^31^. All results in this paper were run with 4 chains, each chain producing 5,000 samples preceded by 5,000 adaptive burn-in samples.

#### MAGeCK

MAGeCK was run using the following parameters:

> mageck test -k [INPUT_COUNTS] -t [LOW_BIN_COLUMNS] -c [HIGH_BIN_columns]

> --sort-criteria pos -n [OUTPUT_DIRECTORY]

Since MAGeCK does not model the bins, we provide the outermost bins in every analysis. Additionally, MAGeCK does not provide a two-sided test at the gene-level. The heuristic we used to deal with this was to take the minimum of the reported gene-level FDR in the positive and negative direction.

#### MAUDE

MAUDE was run using default parameters. For specifics, please see the pipeline code. Since MAUDE requires the gRNA proportions, we gave it the true bin fraction in every simulation. In experimental data we gave MAUDE the GFP+ proportion. Since MAUDE does not have a direct way to merge replicates, we took the gene-level replicate p-values and merged them using Fisher’s method. The resulting Fisher’s p-value was then Benjamini-Hochberg FDR corrected^32^.

## Acknowledgements

We thank Clemens Weiss for helpful comments. This research was supported by NIH grant R01HG008140 and UO1HG012069 (JKP). A.M. is a member of the Parker Institute for Cancer Immunotherapy (PICI), and has received funding from the Innovative Genomics Institute (IGI), the Cancer Research Institute (CRI) Lloyd J. Old STAR award, a gift from the Jordan Family, a gift from the Byers family, and funds from the Simons Foundation and the CRISPR Cures for Cancer Initiative. H.P. is supported by the HHMI Hanna H. Gray Fellowship and the Sloan Foundation Fellowship. Sequencing was carried out at the UCSF CAT, supported by a PBBR grant. We would like to acknowledge the support of the Gladstone Institute Flow Cytometry Core. We would like to thank the Stanford Research Computing Center for providing computational resources and support.

## Author Contributions

Conceptualization, H.P., J.W.F, J.K.P, and A.M.;

Formal Analysis, H.P. and J.W.F.;

Investigation, J.W.F., M.M.A, and C.M.G.;

Resources, J.K.P, and A.M.;

Writing - Original Draft, J.W.F;

Writing - Review & Editing, H.P., J.W.F, M.M.A, J.K.P, and A.M.;

Visualization, H.P. and J.W.F;

Supervision, J.K.P, and A.M.;

Funding Acquisition, J.K.P, and A.M.;

