## Supplementary materials for "A model for accurate quantification of CRISPR effects in pooled FACS screens"

[1] *Department of Genetics, Stanford University, Stanford, CA 94305, USA.*
[2] *Gladstone-UCSF Institute of Genomic Immunology, San Francisco, CA 94158, USA.*
[3] *Department of Microbiology and Immunology, University of California, San Francisco, San Francisco, CA 94143, USA.*
[4] *Department of Medicine, University of California San Francisco, San Francisco, CA 94143, USA.*
[5] *Innovative Genomics Institute, University of California, Berkeley, Berkeley, CA 94720, USA.*
[6] *UCSF Helen Diller Family Comprehensive Cancer Center, University of California, San Francisco, San Francisco, CA 94158, USA.*
[7] *Parker Institute for Cancer Immunotherapy, University of California San Francisco, San Francisco, CA, USA.*
[8] *Institute for Human Genetics, University of California San Francisco, San Francisco, CA 94143, USA.*
[9] *Department of Biology, Stanford University, Stanford, CA 94305, USA.*
[10] *Present address: Genentech Research and Early Development, South San Francisco, CA*
[11] *Departments of Computational Medicine and Human Genetics, University of California, Los Angeles, Howard Hughes Medical Institute, Los Angeles, CA 90024, USA.*
[12] *These authors contributed equally.*
[13] *These authors jointly supervised this work.*

#### 1 The Waterbear model

The Waterbear model has a more general generative view which we use for the simulations and call the "cell-level" view of the model. It is described in Supplementary Section 1.1. The model used for inference is referred to as the "gene-level" model and a full description of the model is in Supplementary Section 1.2.

##### 1.1 The cell-level generative model

The cell-level model can be seen in Algorithm 1. In this model, we assume the gene-level effect sizes, guide-composition (the relative frequency of each guide in the input population), the experiment-level multiplicity of infection (MOI), and bin-sizes are fixed. While this is not necessary, it enables us to define the parameters for these arguments and thus iterate over them, while still keeping randomness in the remainder of the model. Fixed here is also relative to when the model is run. Parameters are fixed at the start of a simulation, but randomly sampled and passed in as arguments in practice. Additionally, a plate model can be seen in Supplementary Figure 1.1.

**Algorithm 1** The *cell-level* generative view of the Waterbear model.

**Input**

$\mu$, a vector of effect sizes, length number of genes. Genes with no effects have value 0.
$\phi$, a vector of length $N$ dispersion values (one per sample).
$\lambda$, the multiplicity of infection.
guide composition, a unit simplex of length number of guides.

**Output**

A tensor of dimension $N$ x number of guides x number of bins.

```
for n in N samples do
    for c in C_n cells do
        M_nc ~ Poisson(λ)                                   ▷ The number of guides in this cell.
        G_nc | M_nc ~ Multinomial(M_nc, guide composition)  ▷ Choose which guides.
        R_nc | G_nc ~ Normal(G_nc^T μ, 1)                   ▷ Given effect sizes μ, draw from the shifted distribution.
        Y_nc ← binMapping(R_nc)                             ▷ Given the reporter, sort the cell into the correct bin.
        B_n[𝟙{G_nc}] ← B_n[𝟙{G_nc}] + Y_nc                  ▷ Update the true cell-guide counts.
    end for
    for g in G guides do
        O_ng ~ DirichletMultinomial(sum(B_n[g]), φ_n B_n[g]/sum(B_n[g]))   ▷ The observation is a noisy version of the truth, post sorting.
    end for
end for
```

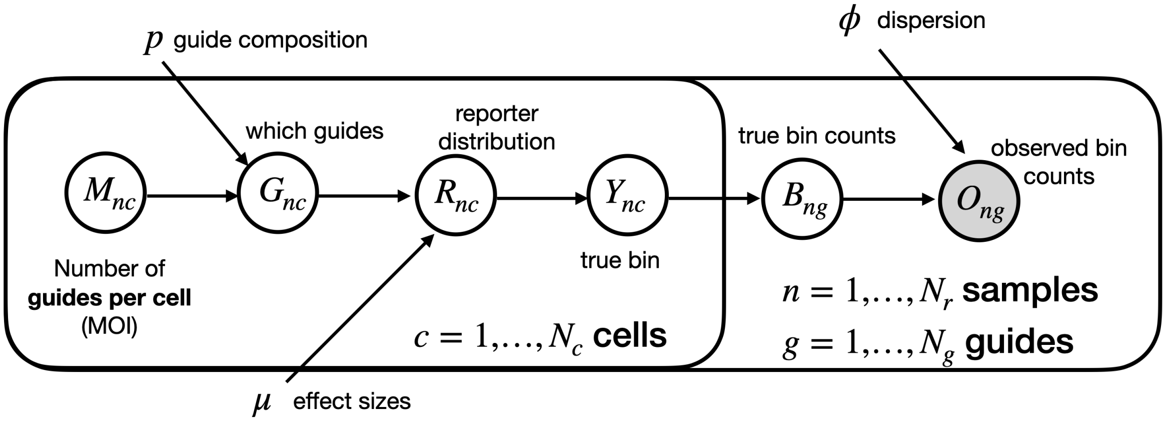


S. Figure 1.1: Plate model for the cell-level model which is used for simulation.

#### 1.2 The gene-level inference model

We begin by describing the full generative model. This model is represented in the plate model in Supplementary Figure 1.2.

First, the experiment-level parameters:

$$\pi \sim \text{Beta}(10, 10),$$
$$\sigma \sim \text{Gamma}(1, 0.10),$$
$$\tau \sim \text{Gamma}(1, 0.10),$$
$$\phi \sim \text{Exponential}(1/\phi_{\text{MLE}}).$$

Importantly, these priors are fairly diffuse. Additionally, $\phi_{\text{MLE}}$ is the MLE estimate learned from a frequentist view of this model only on the non-targetting guides.

Next, we cover the effect parameters,

$$\psi_g \mid \pi \sim \text{Bernoulli}(p),$$
$$\mu_g \mid \psi_g, \sigma \sim \text{Normal}(0, \sigma^2),$$
$$\beta_h \mid \mu_{G(h)}, \tau \sim \text{Normal}(\mu_{G(h)} \cdot \psi_g, \tau^2),$$

where $G(h)$ is a map from the guide index to the corresponding gene index.

Finally, we cover the remaining nuisance parameters. Note that one can specify the model in terms of either the function $\mathbf{q}$ or $t_n$ (see Methods in the main manuscript). Because of this duality, we can specify a prior in either space, and thus choose to do so on $\mathbf{q}$ as the prior is more natural:

$$\mathbf{q_n} \sim \text{Dirichlet}(\alpha),$$
$$\phi_n \mid \phi \sim \text{Exponential}(1/\phi).$$

Here, $\alpha$ is the estimated bin sizes (in unit scale). Finally, we can write the observation as

$$\mathbf{B}_{nh} \mid \mathbf{c}_{nh}, \phi_n, \mathbf{t_n}, \beta_h \sim \text{DirichletMultinomial}(\mathbf{c}_{nh}, \phi_n \mathbf{q}_n(\mathbf{t}_n, \beta_h)).$$

The posterior offers no simple closed form solution, and thus we turn to Markov Chain Monte Carlo. The model is implemented using NIMBLE and can be found on the GitHub link. As implemented, it uses a Gibbs sampler where conditional conjugacy is possible and adaptive random walk MCMC otherwise.

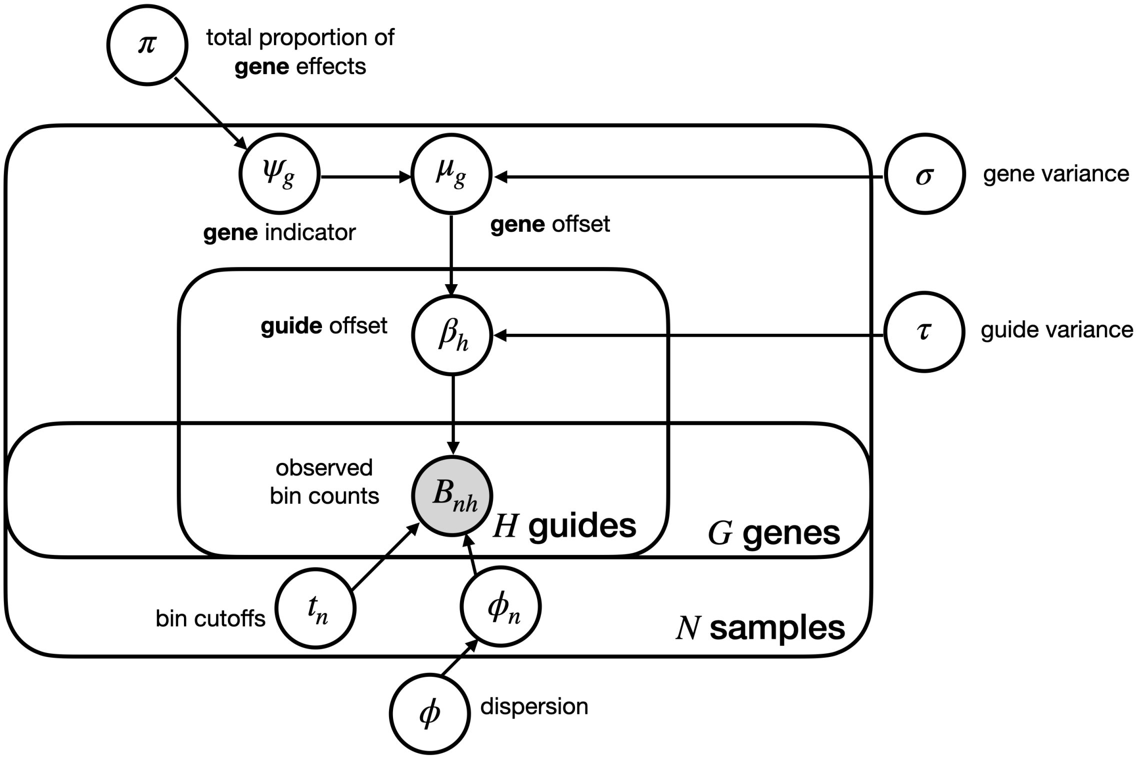


S. Figure 1.2: Plate model for the gene-level model used for inference.

### 2 Simulations

Here, we report additional simulations which were referenced, but not included in the main text for brevity. Additionally, we describe some of the modeling choices (e.g. the effect size generation).

#### 2.1 Simulation effect sizes

Pooled CRISPR screens were performed as described (Methods, Pooled CRISPR screens) and effect sizes were learned from these data. An excess of cells from one donor were collected. After genomic DNA isolation, the genomic DNA was diluted to the equivalent of 50X, 100X, and 200X coverage. Sequencing libraries were made from each diluted sample as described for the other pooled CRISPR screens. We broke the effects into classes easy, moderate, and difficult based on whether MaGeCK called them significant across all dilution rates, some, or only the lowest dilution rate. Since MaGeCK effect sizes are a function of the bin sizes, we used Waterbear to learn the effect sizes in the standard normal space. Those effect sizes are plotted in Supplementary Figure 2.1a. We then smoothed the distribution using the kernel density estimator in R which resulted in the distributions seen in Supplementary Figure 2.1b. Of note, we thresholded the distributions so they would be discrete classes and their intepretation would be simpler. For all simulations, a gene-level effect is drawn from these distributions. To create the *equalmixture* simulations, we sampled classes uniformly at random, then sampled an effect size from the corresponding class. The final distributions are shown in Supplementary Figure 2.2 which shows the marginal distribution of the observed experiments and the simulated experiments is comparable.

#### 2.2 Simulations across additional experimental configurations

In the following simulations we now vary

- the proportion of gene effects (0.05, 0.10, 0.25),
- the number of cells (50e3, 500e3, 1e6),
- the total number of genes (500, 1,000, 5,000),
- the MOI (0.3, 1, 2, 5, 10),

while maintaining four targeting guides per gene assayed, three replicates, and 1,000 non-targeting guides as in the main text. The *proportion of gene effects* is represents the total fraction of genes that have true non-zero effects. The *number of cells* is the number of observed cells across all bins (post FACS sorting). The number of guides is the total number of targeting genes assayed. Each simulation is performed ten times and each line represents an average.

We present the figures grouped by number of genes targeted, as we believe that is most insightful. The figures thus represent 500, 1,000, and 5,000 genes in Supplementary Figures 2.3, 2.4, and 2.5, respectively.

The following overall trends hold true for all figures:

- As the proportion of gene effects increase, sensitivity increases.
- As the MOI increases, sensitivity increases.
- As the number of cells increases, sensitivity increases.

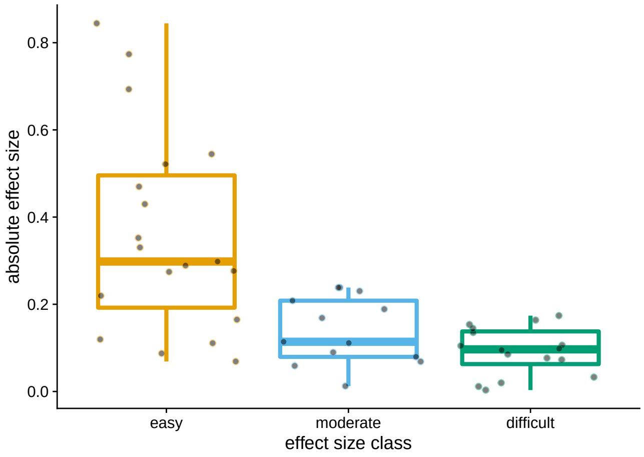


(a) The effect size distribution as inferred from the dilution experiments.

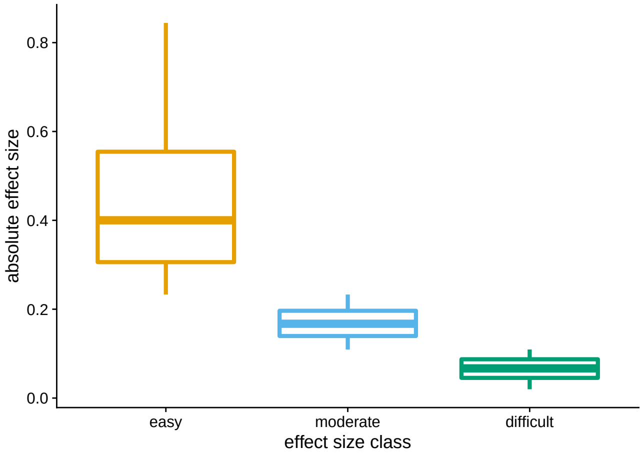


(b) The effect size distribution after smoothing the density.

S. Figure 2.1: Effect size distributions from real data used to inform simulated data.

However, it is clear that this rate reaches an inflection point at 5,000 genes where there is very little sensitivity achievable with 50,000 cells (Supplementary Figure 2.5). Even at 500,000 cells, sensitivity only becomes comparable to the other configurations around MOI 2.

#### 2.3 Effect of Non-Targeting Guides

To test the robustness of inference with varying number of non-targetting guides, we simulated under the similar conditions as in Figure 3 in the main text, while varying only the number of non-targetting guides. In particular, we used 10, 100, and 1,000 non-targetting guides (the main paper uses 1,000). We also maintained the total number of targetting guides. This means that the per guide coverage is technically slightly higher for simulations with smaller number of non-targgeting guides. Supplementary Figures 2.6 and 2.7 display the sensitivity and FDR, respectively. Note, there is little difference, if any in performance as a function of the number of non-targetting guides.

#### 2.4 Effect of unequal bin sizes

Here, we simulated from different bin sizes for each replicate to see how it might change the observed FDR and sensitivity, Supplementary Figures 2.8a and 2.8b. These simulations follow the same sampling scheme as those in Figure 3, with only the bin sizes changed to:

- Replicate 1 bin sizes: (0.10, 0.40, 0.25, 0.25),
- Replicate 2 bin sizes: (0.20, 0.30, 0.30, 0.20),
- Replicate 3 bin sizes: (0.25, 0.25, 0.40, 0.10).

#### 2.5 Sensitivity plots including MAUDE

In the main text we do not include MAUDE in the sensitivity analysis (Figure 3) as it is very miss-calibrated and thus is expected to have high sensitivity somewhat spuriously. Indeed, we see higher sensitivity in Supplementary Figures 2.9a and 2.9b, but perhaps not as high as expected given average FDR around 0.50.

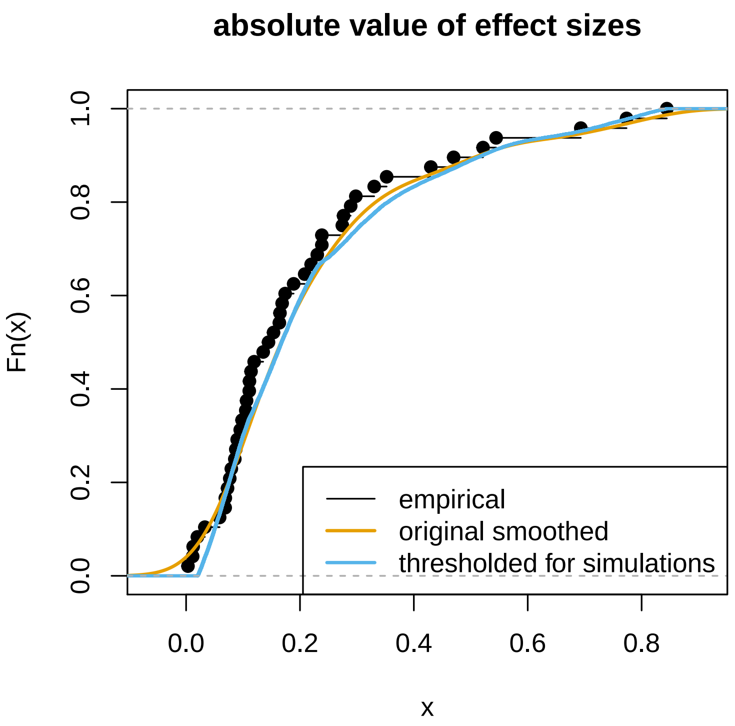


S. Figure 2.2: The ECDF of the complete effect size distribution after pooling easy, moderate, and difficult effects.

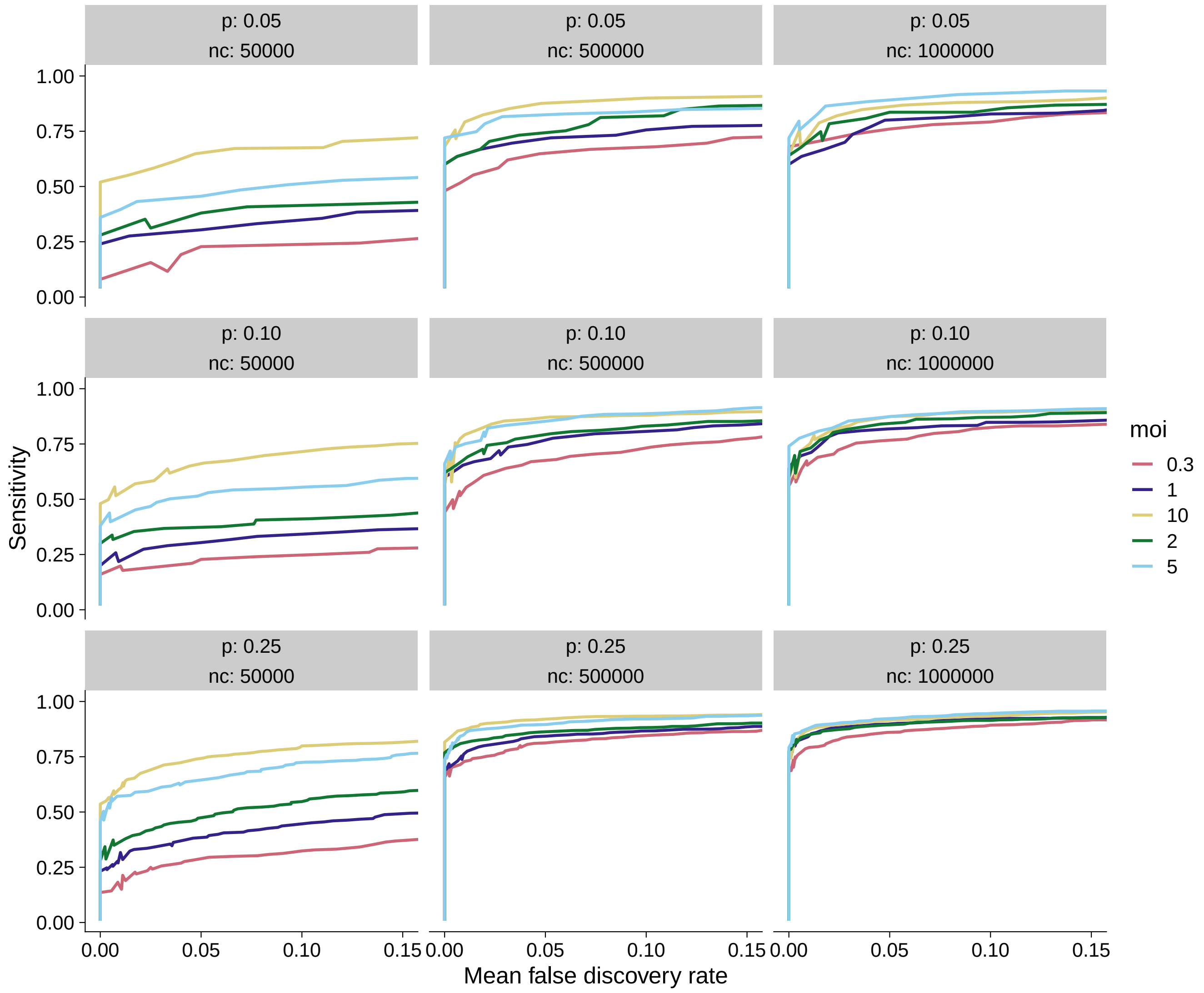


S. Figure 2.3: 500 gene simulations which vary the MOI, proportion of gene effects, and number of cells. In the sub panels, $p$ refers to the proportion of effects, and $nc$ refers to the number of cells.

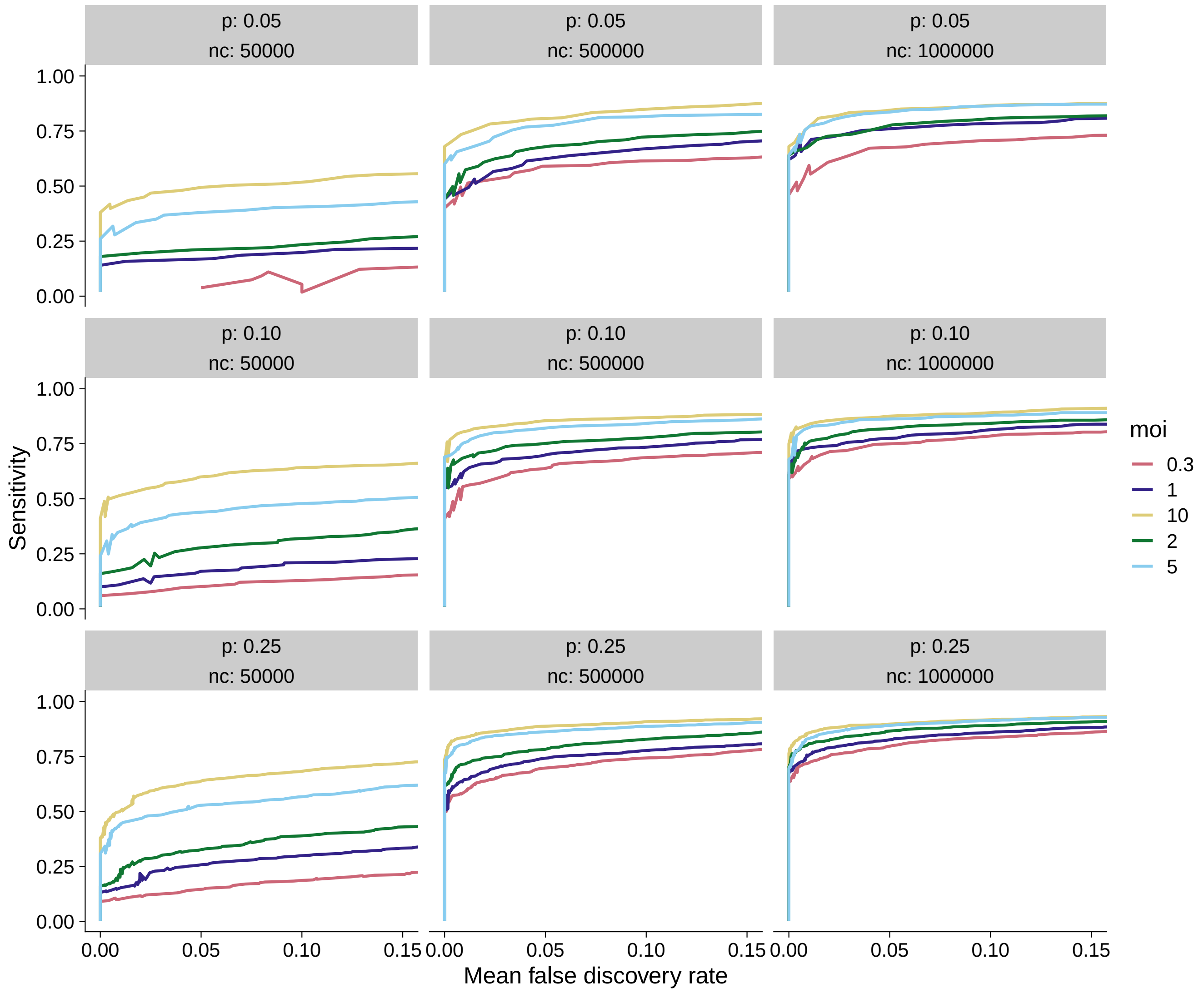


S. Figure 2.4: 1,000 gene simulations which vary the MOI, proportion of gene effects, and number of cells. In the sub panels, $p$ refers to the proportion of effects, and $nc$ refers to the number of cells.

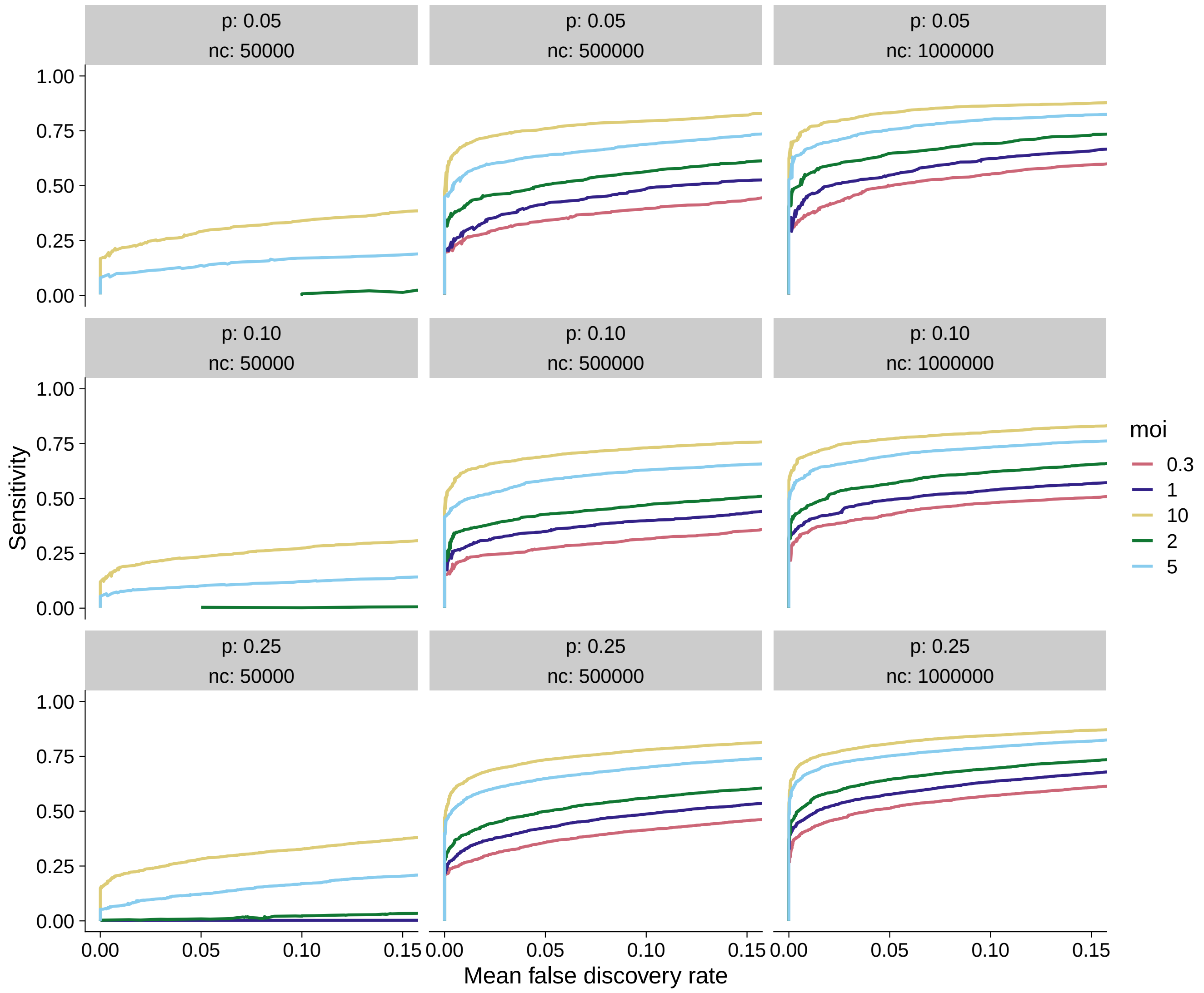


S. Figure 2.5: 5,000 gene simulations which vary the MOI, proportion of gene effects, and number of cells. In the sub panels, $p$ refers to the proportion of effects, and $nc$ refers to the number of cells.

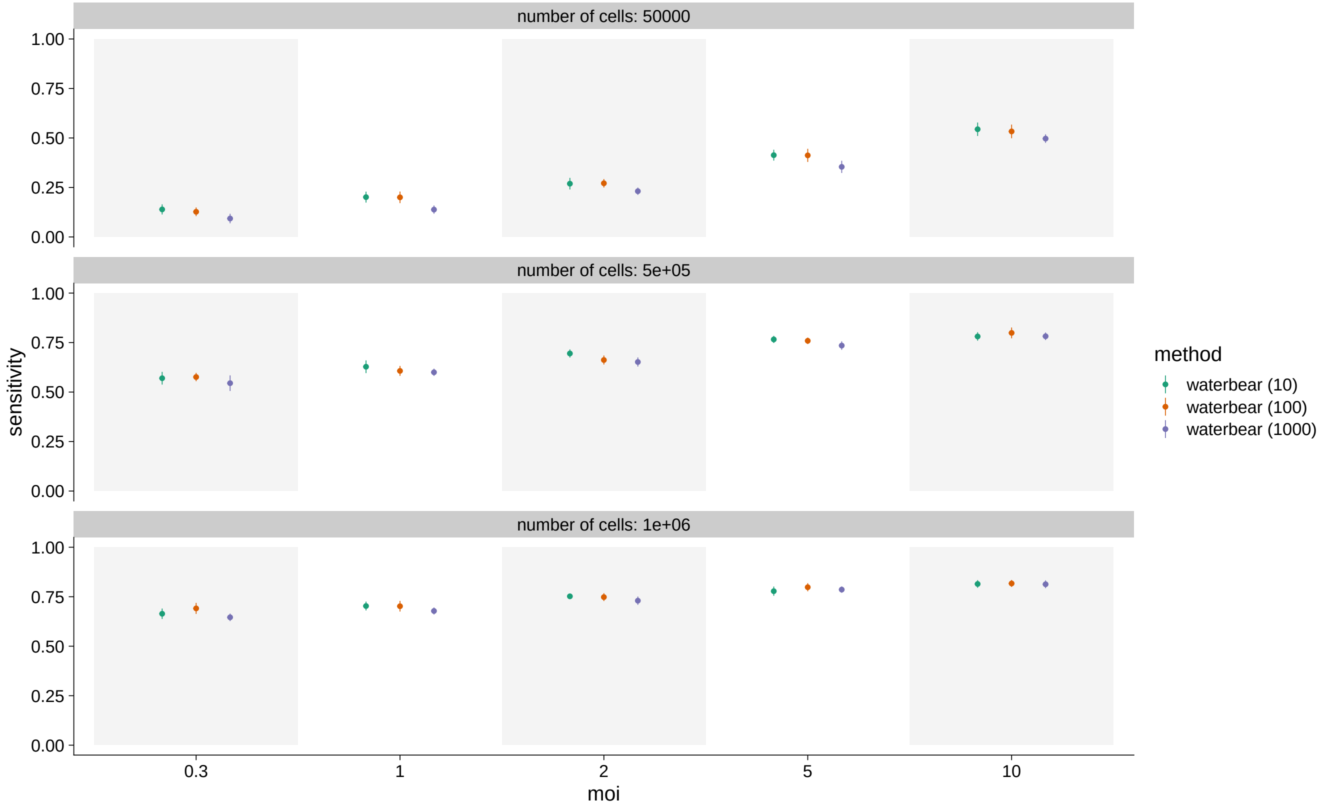


S. Figure 2.6: Sensitivity of Waterbear as a function of number of non-targetting controls (in parentheses), MOI, and number of cells.

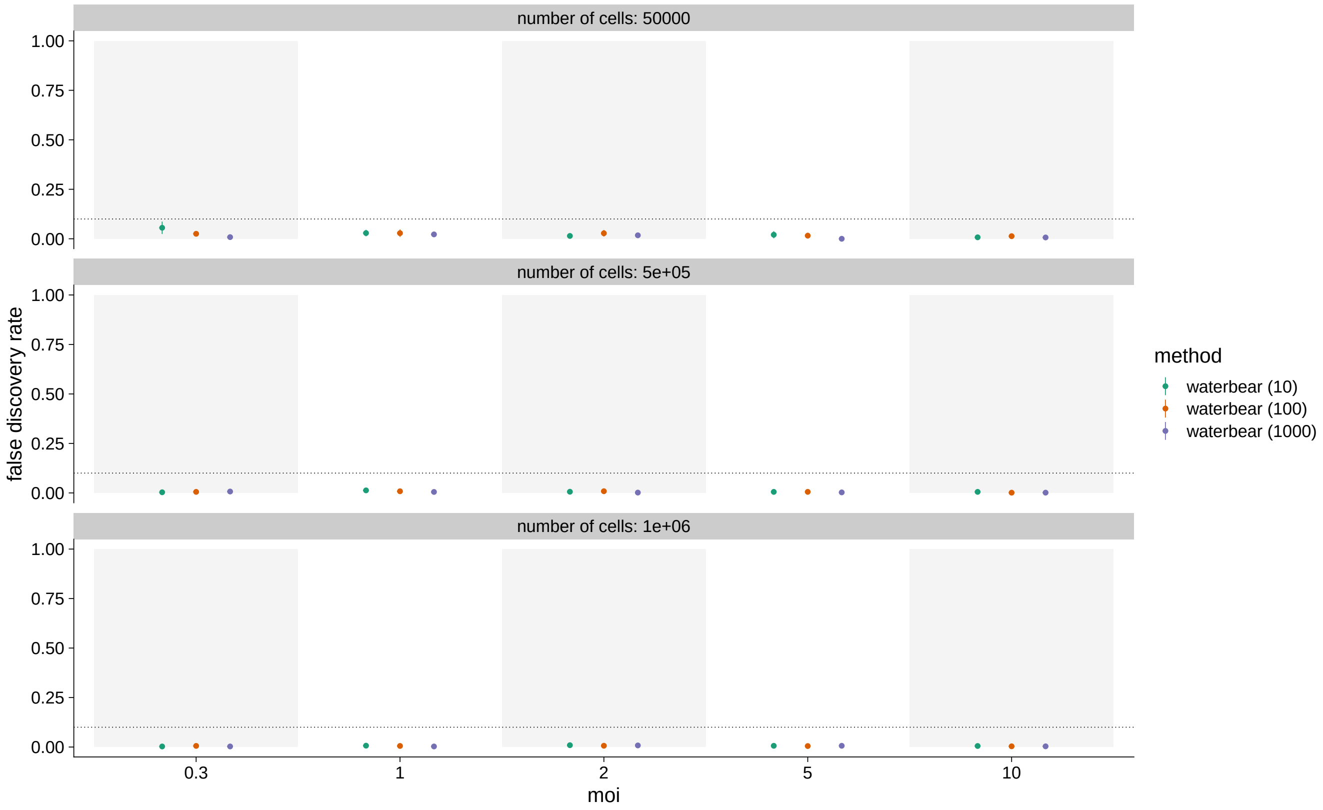


S. Figure 2.7: False Discovery Rate of Waterbear as a function of number of non-targetting controls (in parentheses), MOI, and number of cells.

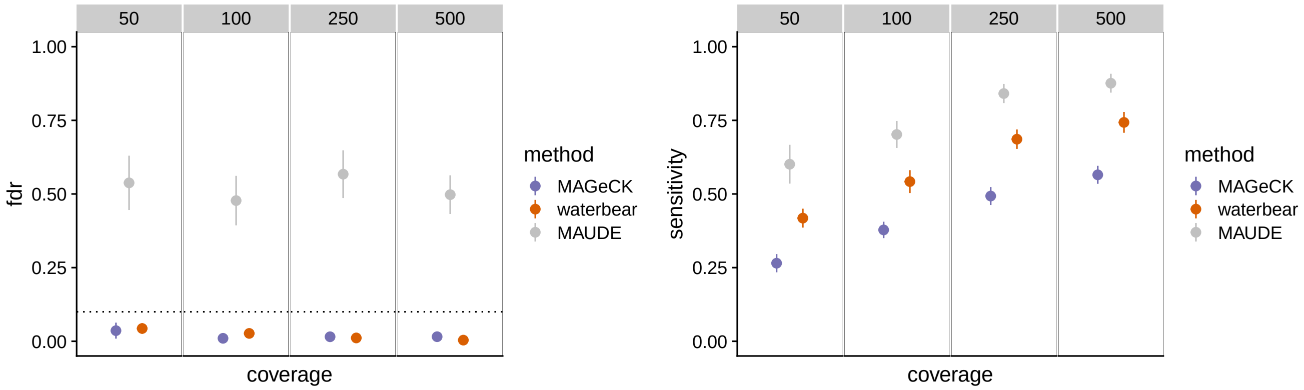


(a) Average FDR results when bin sizes are unequal.

(b) Average sensitivity results when bin sizes are unequal.

S. Figure 2.8: Results from simulations with unequal bin sizes while varying coverage.

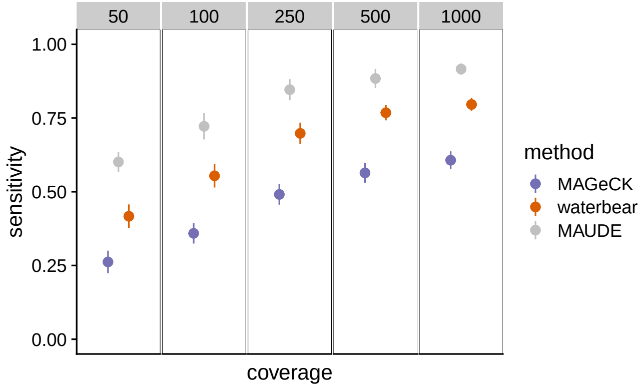


(a) Average sensitivity results while varying coverage and exactly one guide per cell.

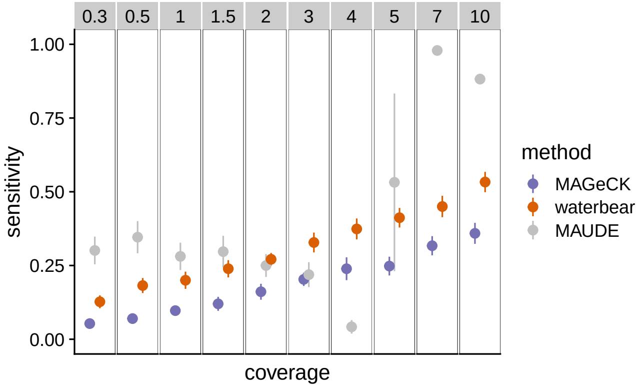


(b) Average sensitivity results while varying MOI.

S. Figure 2.9: Sensitivity results from Figure 3 (main text) including MAUDE.

### 3 Experimental validation results

In this section we display further results from the arrayed knockdown experiments [1]. We previously validated a number of regulators of IL2RA by individually knocking out each regulator and measuring the effect on IL2RA levels (Freimer et al.). Here we use that data to validate Waterbear's sensitivity identifying regulators from IL2RA CRISPR FACS screens under different experimental conditions including changing MOI, coverage, and number of replicates.

#### 3.1 Down sampling replicates

Here, our goal is to see how much the results are affected by reducing the number of replicates from three to two. Of particular interest is the extreme of high-coverage, low MOI (a conventional experiment) to low-coverage, high MOI. These figures can be seen in Supplementary Figures 3.1 - 3.4. In particular, we note reasonable concordance between all four cases, however, we note that (low coverage, low MOI) has the lowest, while (high coverage, high MOI) has the highest concordance.

#### 3.2 Validation against arrayed knockouts

Here, we use the arrayed knockout data from [1] to benchmark all the tools against a positive control set (26 genes) and a much smaller null set (2 genes). These genes are listed in Supplementary Table 1. Waterbear is shown here with all gene names (Supplementary Figure 3.5), along with MaGeck (Supplementary Figure 3.6), and MAUDE (Supplementary Figure 3.7). In particular, both Waterbear and MAUDE show high sensitivity, whereas MaGeCK misses a large number of true positives, even some with modest effect sizes (e.g. JAK3, RELA).

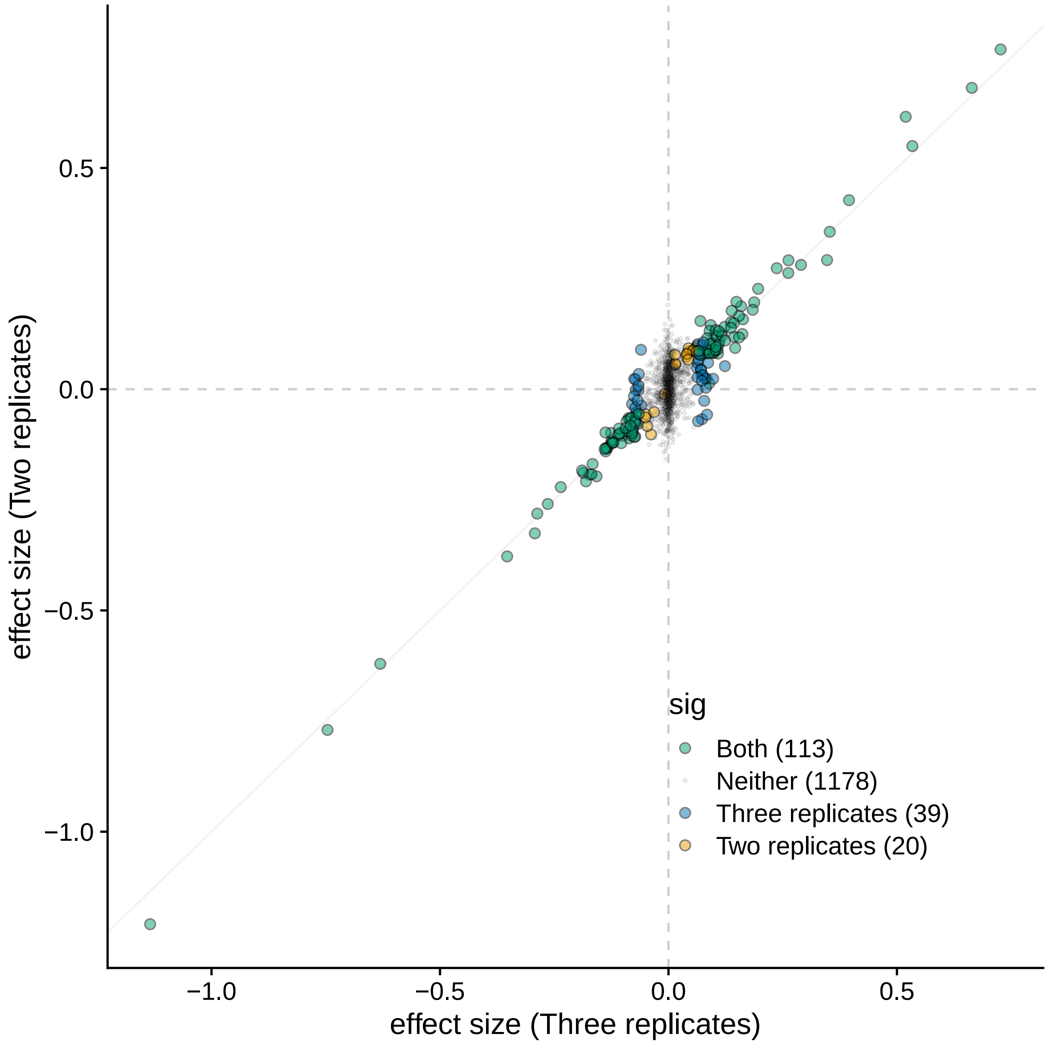


S. Figure 3.1: The effect size distribution and significant hits at LFSR $\leq 0.10$ when including only replicates 1 and 2 versus all three replicates with the high coverage, low MOI data.

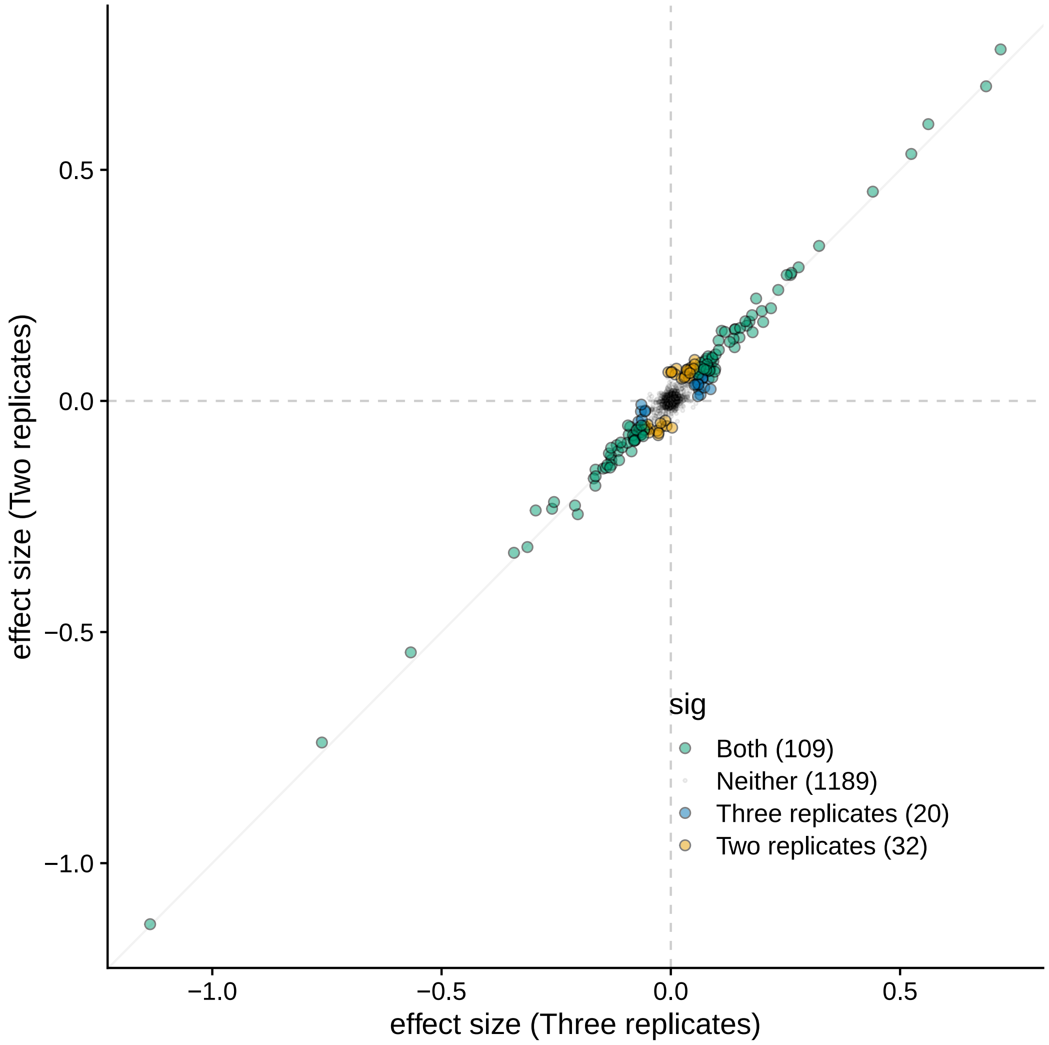


S. Figure 3.2: The effect size distribution and significant hits at LFSR $\leq 0.10$ when including only replicates 1 and 2 versus all three replicates with the high coverage, high MOI data.

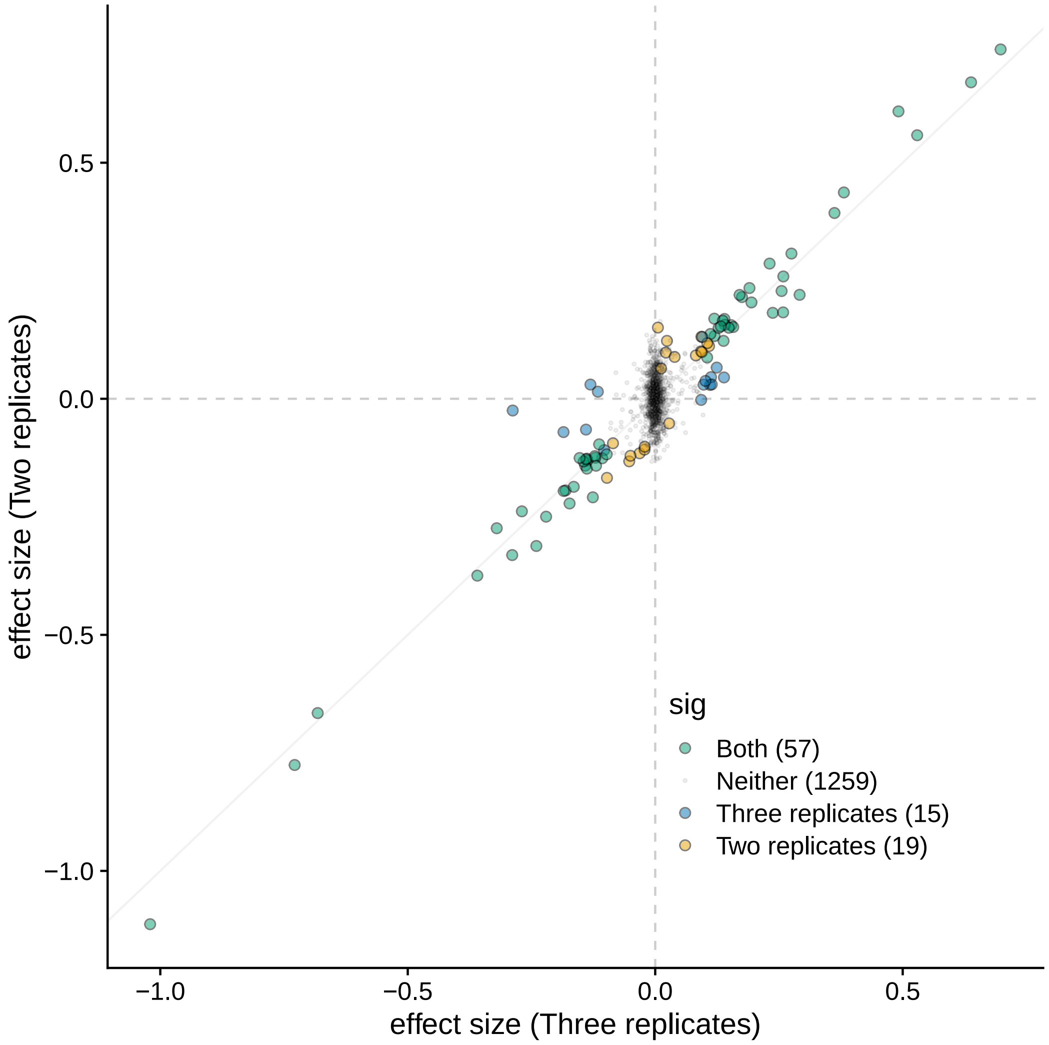


S. Figure 3.3: The effect size distribution and significant hits at LFSR $\leq 0.10$ when including only replicates 1 and 2 versus all three replicates with the low coverage, low MOI data.

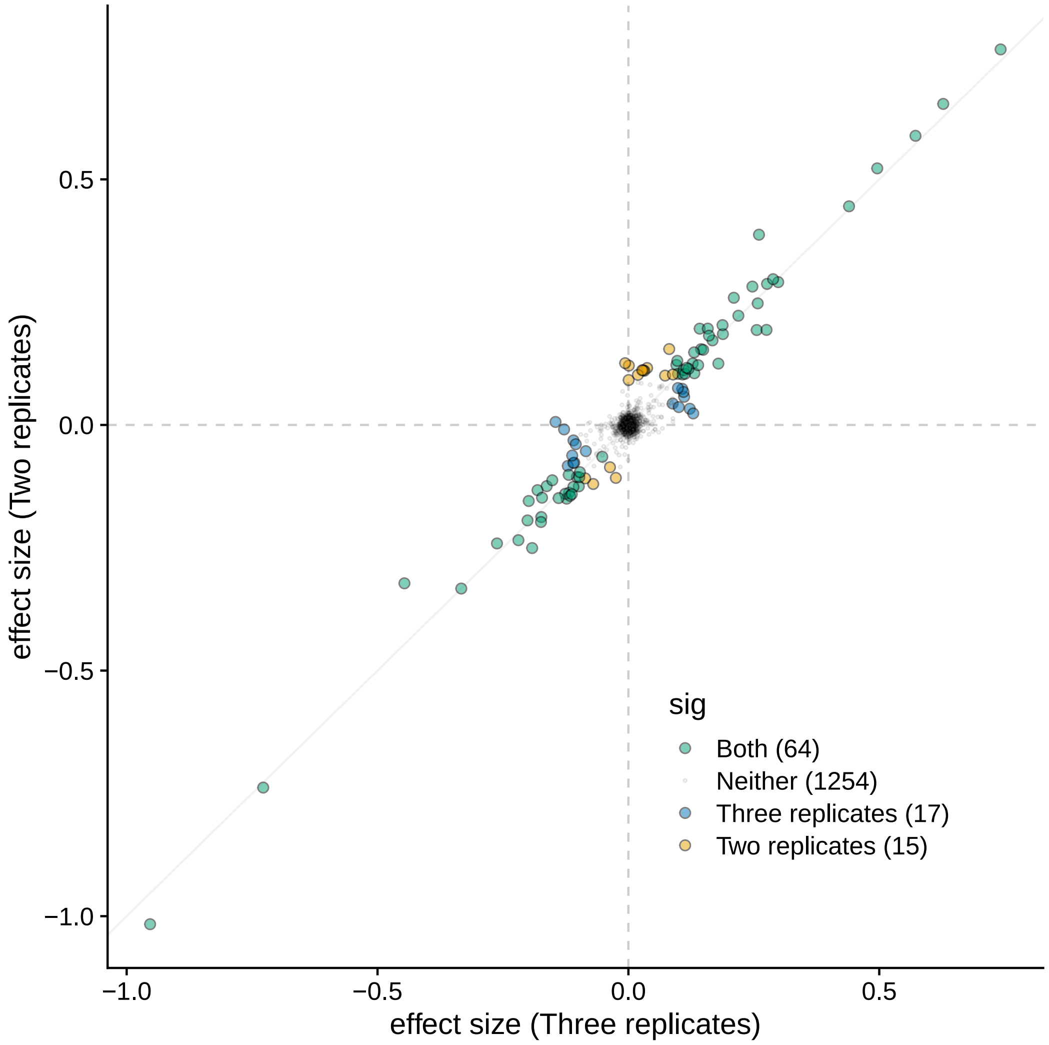


S. Figure 3.4: The effect size distribution and significant hits at LFSR $\leq 0.10$ when including only replicates 1 and 2 versus all three replicates with the low coverage, high MOI data.

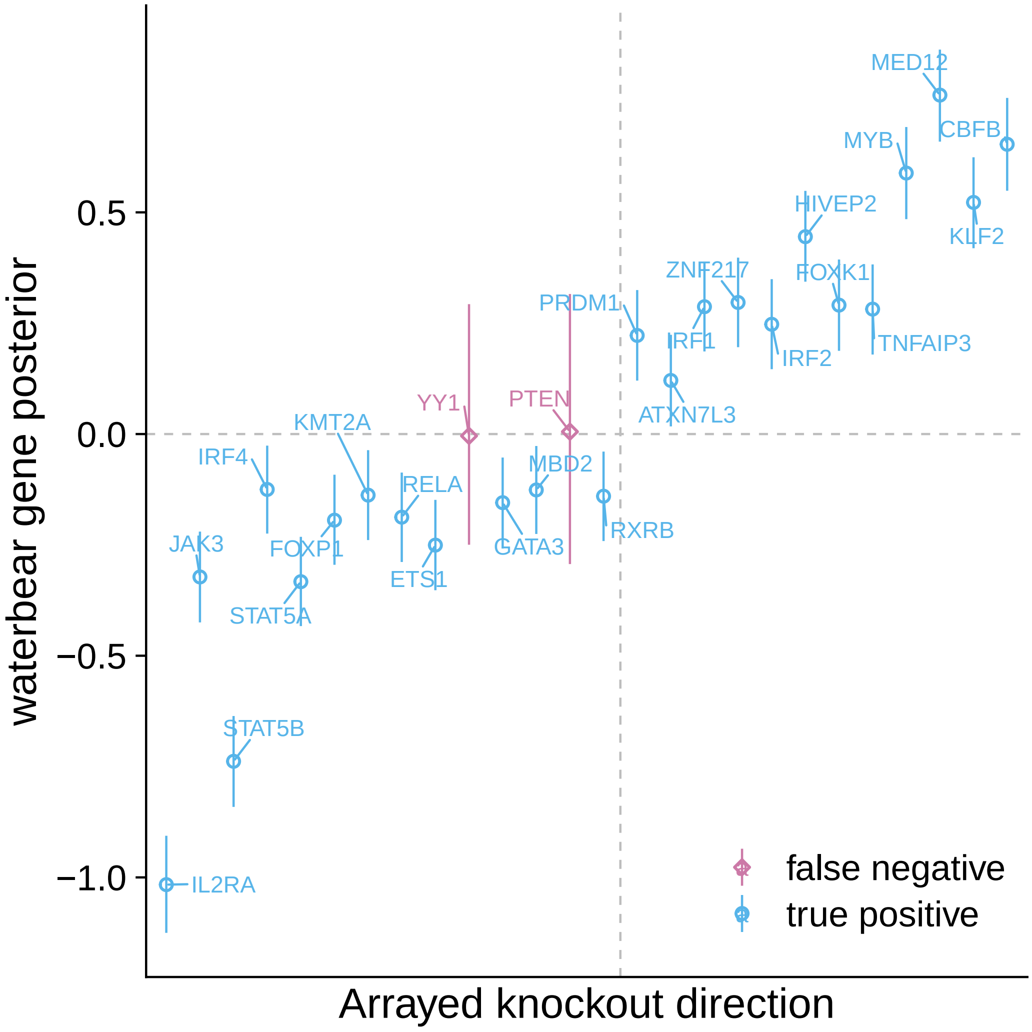


S. Figure 3.5: Validation using low coverage, high MOI data for Waterbear.

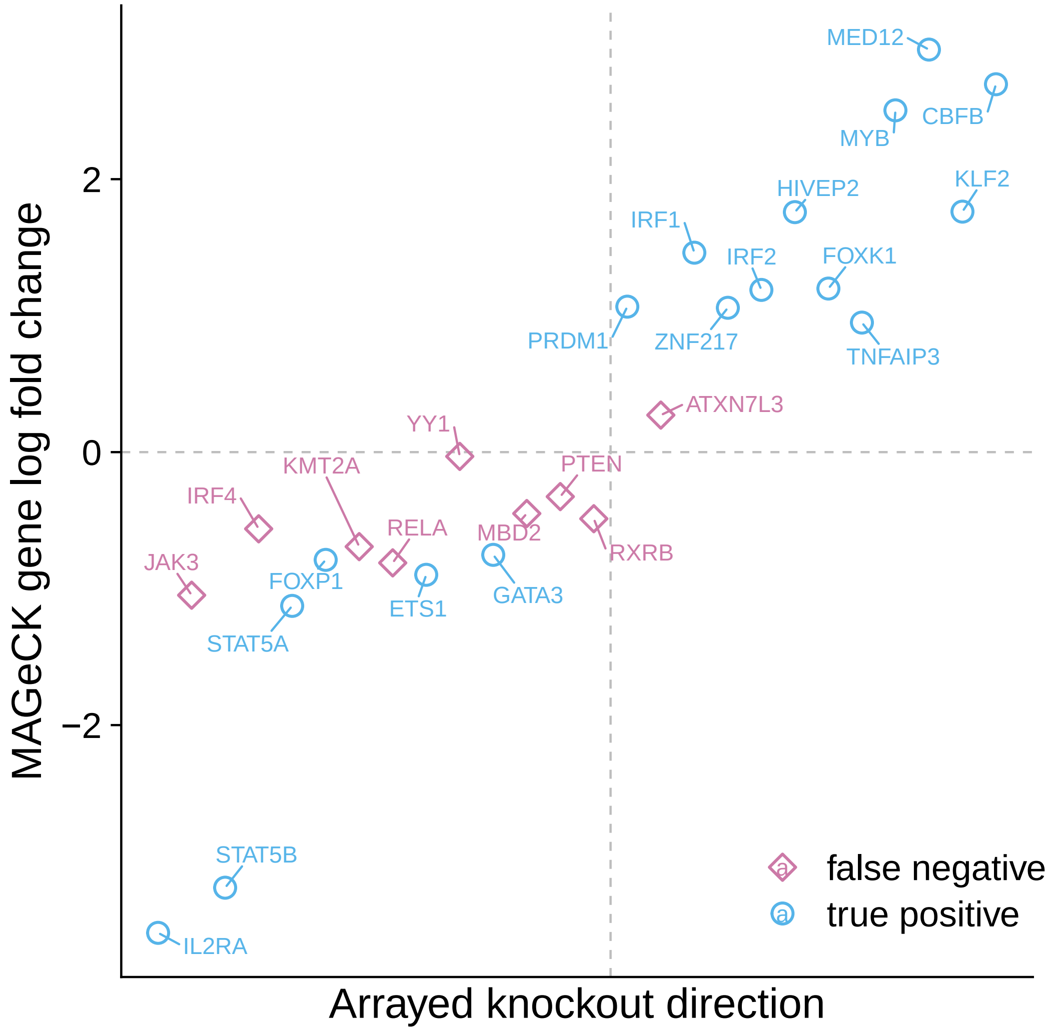


S. Figure 3.6: Validation using low coverage, high MOI data for MaGeCK.

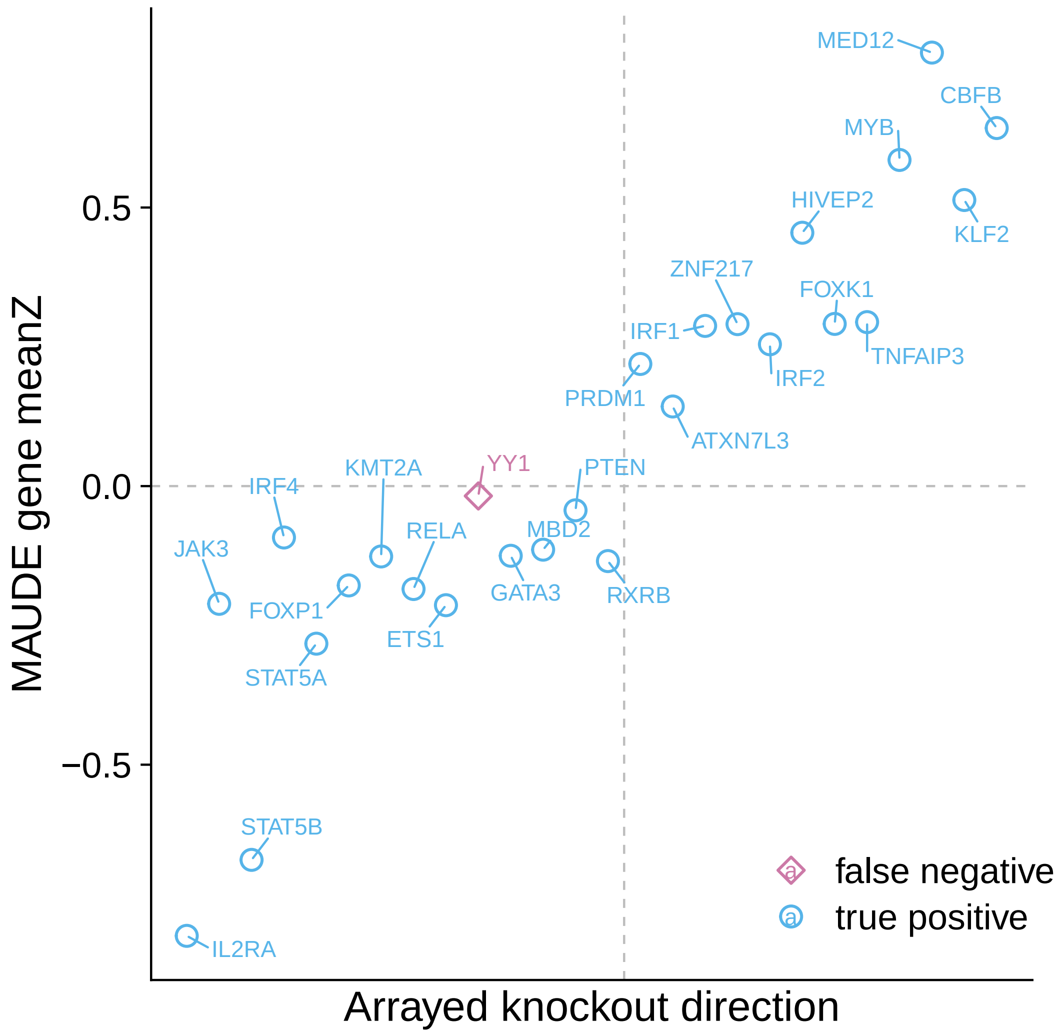


S. Figure 3.7: Validation using low coverage, high MOI data for MAUDE.

| | gene | validation_direction | mfi_avg | mfi_rank | mfi_rank_centered |
|---|---|---|---|---|---|
| 1 | ETS1 | Decrease | -0.53 | 9.00 | -5.50 |
| 2 | ATXN7L3 | Increase | 0.30 | 16.00 | 1.50 |
| 3 | MYB | Increase | 0.79 | 23.00 | 8.50 |
| 4 | KMT2A | Decrease | -0.68 | 7.00 | -7.50 |
| 5 | TNFAIP3 | Increase | 0.66 | 22.00 | 7.50 |
| 6 | GATA3 | Decrease | -0.43 | 11.00 | -3.50 |
| 7 | STAT5B | Decrease | -2.40 | 3.00 | -11.50 |
| 8 | YY1 | Decrease | -0.44 | 10.00 | -4.50 |
| 9 | FOXP1 | Decrease | -0.75 | 6.00 | -8.50 |
| 10 | FOXK1 | Increase | 0.59 | 21.00 | 6.50 |
| 11 | STAT5A | Decrease | -1.54 | 5.00 | -9.50 |
| 12 | RXRB | Decrease | -0.23 | 14.00 | -0.50 |
| 13 | MBD2 | Decrease | -0.38 | 12.00 | -2.50 |
| 14 | MED12 | Increase | 0.87 | 24.00 | 9.50 |
| 15 | ZNF217 | Increase | 0.37 | 18.00 | 3.50 |
| 16 | KLF2 | Increase | 0.92 | 25.00 | 10.50 |
| 17 | IRF1 | Increase | 0.32 | 17.00 | 2.50 |
| 18 | HIVEP2 | Increase | 0.45 | 20.00 | 5.50 |
| 19 | PRDM1 | Increase | 0.29 | 15.00 | 0.50 |
| 20 | PTEN | Decrease | -0.35 | 13.00 | -1.50 |
| 21 | IRF2 | Increase | 0.38 | 19.00 | 4.50 |
| 22 | RELA | Decrease | -0.68 | 8.00 | -6.50 |
| 23 | CBFB | Increase | 1.62 | 26.00 | 11.50 |
| 24 | IL2RA | Decrease | -4.80 | 1.00 | -13.50 |
| 25 | IRF4 | Decrease | -2.26 | 4.00 | -10.50 |
| 26 | JAK3 | Decrease | -3.05 | 2.00 | -12.50 |

Table 1: The validation set read from [1]. Genes that were not included due to uncertain effects were: MYC, IKZF1, IKZF3, FOXO1, and SMARCB1.

### 4 Average number of collisions

Here, we derive a simple model to estimate the number of *collisions*. A *collision* occurs when two or more guides which have an effect on the phenotype enter the same cell. By definition, collisions are an upper bound of the expected fraction of epistatic events, since two guides may enter a cell and have independent effects on the phenotype. Our goal here is to compute for a given cell, the expectation that this cell receives two or more guides with an effect. In particular, there are two free parameters in this calculation: (1) $p$, the proportion of guides that have an effect on the phenotype, and (2) $\lambda$, the multiplicity of infection.

Let $X$ be a random variable that counts the number of guides in a cell. We assume,

$$X \sim \text{Poisson}(\lambda),$$

where $\lambda$ is the multiplicity of infection (MOI) of the experiment. If we assume that $p$ proportion of total guides in our pool have an effect, then, if we select a guide at random, let

$$G_i = \begin{cases} 1 & \text{if guide } i \text{ has an effect,} \\ 0 & \text{otherwise.} \end{cases}$$

Further, we assume $P(G_i = 1) = p$ (a Bernoulli random variable). Finally, define,

$$Y = \mathbb{1}\left\{ \sum_{i=1}^{X} G_i \neq 0 \right\} = \mathbb{1}\left\{ \sum_{i=1}^{X} G_i > 0 \right\}.$$

By definition, $Y$ is 1 if a collision happens, and 0 otherwise. Thus, by construction, we want to compute $E[Y] =$

$P(\sum_{i=1}^{X} G_i > 0)$, the probability of a collision. By iterated expectation we get,

$$E[Y] = E_X\left[E[Y \mid X]\right] = E_X\left[P\left(\sum_{i=1}^{X} G_i > 0 \mid X\right)\right].$$

Additionally, note that

$$\begin{aligned} P\left(\sum_{i=1}^{X} G_i > 0 \mid X\right) &= 1 - P\left(\sum_{i=1}^{X} G_i = 0\right) \\ &= 1 - P(G_1 = 0, \ldots, G_X = 0 \mid X) \\ &= 1 - (1-p)^X. \end{aligned}$$

This result yields,

$$\begin{aligned} E[Y] &= E_X[1 - (1-p)^X] \\ &= 1 - E_X\left[(1-p)^X\right]. \end{aligned}$$

Note that, the moment generating function of $X$, a Poisson, is

$$E[\exp(tX)] = \exp(\lambda(\exp(t) - 1)).$$

By taking $t = \log(1-p)$,

$$E[\exp(\log(1-p)X)] = E_X[(1-p)^X].$$

Thus, the expected number of collisions per cell is a function of the multiplicity of infection, $\lambda$, and the proportion of guides that have an effect, $p$,

$$\begin{aligned} E[Y] &= 1 - \exp\left[\lambda(\exp(\log(1-p)) - 1)\right] \\ &= 1 - \exp\left[\lambda(1 - p - 1)\right] \\ &\boxed{= 1 - \exp(-\lambda p).} \end{aligned}$$

This function can be visualized in Supplementary Figure 4.1.

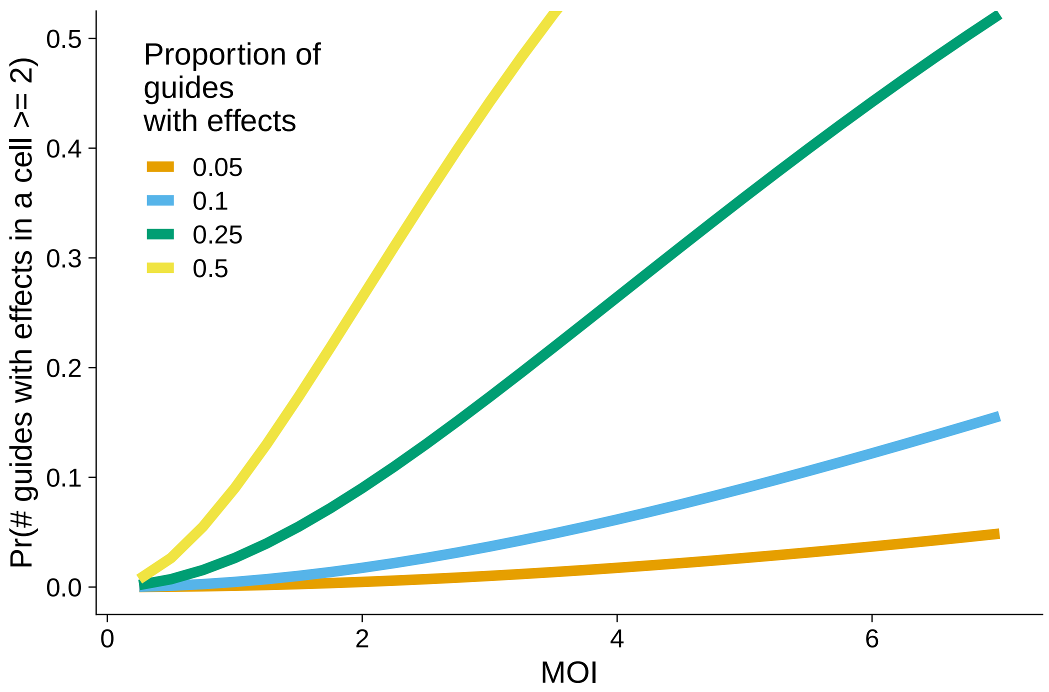


S. Figure 4.1: The average expected number of collisions as computed in Supplementary Section 4.
